# Polyploid *Achromatium* sp. expresses protein variants based on environmental cues

**DOI:** 10.64898/2026.09.07.749883

**Authors:** Sophie-Luise Heidig, Sunu Lama, Jeewan-Babu Rijal, Pierre H. Remy, Jean-Marie Volland, Olivier Gros, Wim F. Vranken, Christine Carapito, Danny Ionescu

## Abstract

*Achromatium* spp. are giant bacteria harboring hundreds of non-clonal chromosomes per cell. The allelic divergence they exhibit could be a mechanism to enable fine-tuned responses to environmental changes, enabling this organism to broaden its ecological range. However, while ubiquitous in freshwater and marine environments, *Achromatium* has not been obtained in culture, leaving the role and functioning of its genomic and proteomic diversity poorly understood. Here, we incubated freshly collected freshwater *Achromatium* under three temperature conditions to conduct comparative mass spectrometry-based proteomics experiments. These were interpreted against an *Achromatium*-specific protein atlas developed from curated published and new assemblies from marine and freshwater environments. This *de novo* assembled atlas allowed a twenty-fold increase in the number of identified proteins compared to mass spectrometry data searches against the UniProt database. Predicted proteins from the *Achromatium* atlas were grouped into orthologs, 5 % of which showed temperature-associated detection of distinct protein variants within the group. This provides the first indication that *Achromatium* uses distinct protein variants under different environmental conditions, enabling it to swiftly respond to environmental changes through differential expression from its cache of chromosomes.

## Main

Most bacterial cells are treated as genetically uniform units, but *Achromatium* challenges this view. A single *Achromatium* cell can contain approximately 150–400 non-clonal chromosome copies, with reported sequence identities ranging from 80 % to 99 %^1^. Thus, an individual *Achromatium* cell inherently contains a diverse repertoire of related genes. This raises a central question on the ecological role of chromosome-level diversity. Does it merely provide genetic redundancy, or can it serve as a functional reservoir from which different protein variants are expressed under different environmental conditions? If such variants are expressed selectively, a single cell could adjust its physiology by drawing on its existing chromosome repertoire rather than relying on new mutations. Such behavior has been observed in eukaryotes before, specifically in the temperature adaptation of the diatom *Fragilariopsis cylindrus*^2^. This could help explain how *Achromatium* survives globally in marine, brackish and freshwater ecosystems without apparent significant differences in its functional repertoire across different environment types^3^.

Cells of *Achromatium* can exceed 100 μm in length^1,3^, making it the largest known freshwater bacteria. Even larger bacteria with even more chromosomes have been found in marine environments. An example is *Thiomargarita magnifica*, a giant bacterium reaching up to 1 cm in length and containing hundreds of thousands of largely clonal chromosomes^4,5^. Why some polyploid giant bacteria contain more chromosomal diversity than others remains unresolved^4^. Since *Achromatium’s* chromosomal heterogeneity is well documented^1,3,5–8^, it offers an opportunity to investigate whether polyploidy can also support functional diversity within a single cell. Defined in prokaryotes as having more than 10 chromosomes per cell, polyploidy has been described in a growing number of bacterial species in a temporary or permanent manner, where it may provide several physiological advantages^6,8–12^. In giant bacteria, multiple chromosomes help to compensate for limited intracellular transport and reduced diffusion efficiency by spatially coupling transcription and translation to substrate-containing regions^5^. Gene redundancy can also buffer the effects of deleterious mutations while allowing novel functions, traits, and ecological niches to be explored^9^. Multiple genome copies may also increase metabolic capacity through parallel transcription and translation from spatially separated sites^11^.

Mass spectrometry (MS)-based proteomics provides a direct means of determining which elements of the extensive protein repertoire encoded by *Achromatium* are expressed under specific environmental conditions. In polyploid organisms, proteomics provides a key advantage over transcriptomics, as the RNA level is subject to regulatory complexity that can obfuscate the final functional protein machinery of the cell^13^. Direct access to the actual amino acid sequences of the proteins that constitute the *Achromatium* proteome in a given environment then enables interpretation of their general 3D shape, but also their flexibility, the constant movement that governs protein activity^14^. Since protein sequences are shaped by structural and functional constraints and evolve at different rates across sites^15,16^, proteins with related functions can accumulate substantial sequence divergence while retaining evolutionary relationships, thus forming groups of ortholog proteins (OGs)^17^. Within these OGs, the overall 3D fold of the proteins remains generally conserved even when mutations accumulate^18,19^. However, these substitutions can still influence the biophysical properties of the sequences, such as the flexibility or dynamics of the protein backbone, which are more sensitive to environmental conditions^20,21^. Comparing backbone dynamics propensities per position among sequences within the same OG can therefore provide an indirect proxy for the potential structural and functional consequences of sequence variation.

In this study, we combined our analysis of the theoretical proteome derived from genomic data with experimental, MS-based, proteomics to investigate whether the chromosome-level diversity of Achromatium is associated with condition-dependent expression of protein variants and addressed three related questions. First, do the available assemblies contain distinct sequence variants within protein families, rather than divergence generated through random drift? Second, are these variants associated with habitat type, e.g. freshwater versus marine origin? Third, can MS-based proteomics data help determine whether certain protein variants are preferentially expressed following incubation at different temperatures? Temperature provides a useful environmental variable for experimental validation as it can change over relatively short timescales and is known to influence protein stability, metabolism, and cellular stress responses^2^. This approach allowed us to examine whether the unusual chromosome-level diversity of *Achromatium* may contribute to short-term physiological flexibility.

## Results

### Theoretical proteome of *Achromatium*

To obtain an overview of the overall proteomic diversity of *Achromatium* across different environments, we assembled the complete set of proteins encoded in 51 curated assemblies available in July 2025 (Tab. S 1) into the *Achromatium* atlas (see Methods, Data collection and cleaning). We refer to this atlas as the “theoretical proteome” of *Achromatium*. The assemblies originated from freshwater (Lake Stechlin^1,6^: 40, Lake Pavin^22^: 5) and saline environments (Guadeloupe mangroves: 2, Tide pool^23^: 1, Warm mineral springs^7^: 3). From these assemblies we obtained 2259 OGs containing more than 10 genes, with an average of 71 genes (Fig. S1). Initially, functional annotations were available for less than half of the proteins. Combining OGs and annotations we could extend the number of annotated proteins to 70 %. No single copy orthologs were identified. For most OGs, more than one protein copy per assembly was present. OG0000000, the largest OG with 5096 genes, was excluded from the analysis due to the low quality of the multiple sequence alignment (MSA). Many proteins in the OG contain the extracellular peripheral membrane domain DUF1566, also known as Legionella collagen-like domain or IPR011460^24^ (Tab. S 2), as part of their sequence. The internal structure of *Achromatium* is dominated by internal cell membranes that envelope its characteristic calcium carbonate bodies, therefore it stands to reason it developed unique ways to utilize this region, along with novel proteins whose function is yet to be determined.

Nearly half of the OGs harbor distinct variant clusters with changes in biophysical properties rather than random substitutions. The biophysical properties, such as the backbone dynamics, were predicted using the b2bTools^16^ for all OGs. To assess the internal data structure, we calculated a pairwise sequence-identity-based distance matrix using global and pairwise sequence alignment tools, applied k-Means based clustering and evaluated the geometric separation of the clusters using the Silhouette score (SIL). SIL scores range between -1 and 1, with values around 0 representing overlapping clusters and values higher than 0.5 indicating a reasonable separation of clusters. We repeated this clustering with a second distance matrix based on the absolute difference between aligned per-residue predictions of protein backbone flexibility. Within 43 % of OGs, distinct sequence-identity based clusters with changes in biophysical properties exist (SIL>0.5, Fig. S2A). Using global and pairwise alignment methods did not cause a meaningful change (Fig. S2).

Sequence clusters within OGs do not correlate in most cases with external factors such as shared environment (freshwater or marine) (Fig. 1A, C) or shared ancestry (shared sampling location, e.g. Lake Stechlin) (Fig. 1B, D). The external factors clustering was performed by assigning cluster-membership labels based on the sampling information of the assemblies and scored using SIL. We further assessed the match between k-Means-assigned (internal) labels and the external labels using the Adjusted Rand Index (ARI). The ARI scores range between -1 and 1, with values around 0 representing random assignment to clusters values higher than 0.5 indicating a reasonable agreement of internal and external clusters. For most OGs, the relationship between external factors and internal data structure is random, with ARI scores of ±0.1 and SIL scores between -0.2 and 0.1 (Fig. 1). Clustering based on sequence identity (Fig. 1A, B) and backbone dynamics (Fig. 1C, D) capture different aspects of the variant diversity. Applying a threshold of 0.5 for both SIL and ARI, only 7 (Fig. 1A) OGs show a positive relationship for the clustering for environment as compared to 17 for backbone dynamics, and 18 OGs (Fig. 1B) versus 35 OGs (Fig. 1D) show a positive relationship for the clustering for sampling location. Less than 1 % of OGs variant diversity is specific for marine versus freshwater environments considering only sequence identity, with a small increase if biophysical factors are included.

**Fig. 1:**
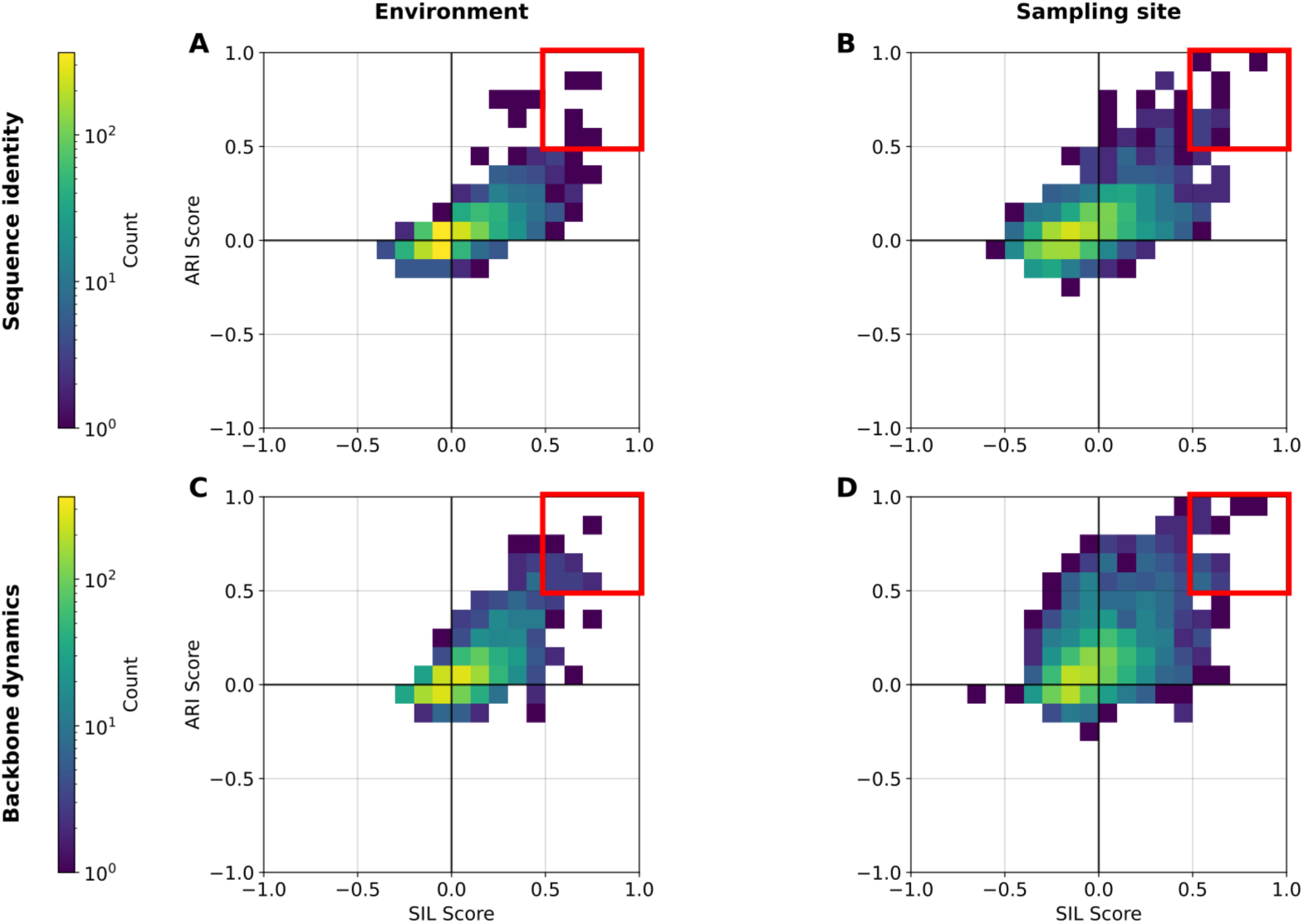
Variant assignment is independent from shared ancestry and environment. External labels mapped from assembly information on marine/freshwater origin (A, C) or sampling location (B, D). Internal clusters calculated based on sequence identity (A, B) or differences in protein backbone dynamics (C, D). The panels compare scoring of external label clustering (SIL) with matching of external and internal clusters (ARI). Score distribution for Environment (A, C) centers mainly around the axis origin, scoring for sampling site (B, D) is more spread out. Scores for ARI and SIL range between -1 (worse than random), 0 (random) to 1 (perfect label matching/geometric separation), with the red box indicating the 0.5 threshold. Number of OGs per bin is shown in color on a logarithmic scale.

### Comparative proteomic analysis

We obtained MS-proteome data for the selective incubation of *Achromatium* at three different temperatures between 7-30 °C (C, M, H) and three time-points up to 4 weeks (T0, T1, T2) (see Methods, Selective incubation). Results were obtained in triplicates for five of the seven investigated conditions. In condition H at T2, only one sample survived (Tab. S 3). Extending the standard UniProt database with the non-redundant *Achromatium* atlas allowed us to increase the number of proteins identified nearly 20-fold from 600 to 17,000 proteins (see Methods, Protein identification and quantification). No proteins were identified in one replicate for condition H at T1. The lowest protein numbers were identified in H samples (Fig. S3). Most proteins were present in both cold and medium conditions (CM) or all conditions (CMH) (Fig. S4). The sequence diversity of *Achromatium* allowed to distinguish individual variants via their unique peptides, thus providing evidence for their condition-specific expression. In a variant-rich organism, such condition-associated detection patterns may reflect selective representation of protein variants in the measured proteome. Differential expression analysis is very sensitive to such intermitted protein identification. The significant contrast would be driven by proteins detected in one condition but not the other, rather than by consistent abundance shifts across biological replicates. We therefore directly use the protein detection patterns as evidence of condition-associated variant expression in the downstream analysis.

The experimental proteomics dataset contained 1456 of the 2259 OGs included in the theoretical proteome of the *Achromatium* atlas. Complementary to this, we assessed the completeness of the recovered proteome from the experiment by mapping the detected proteins to the *Chromatiaceae* set of KEGG proteins^25^ (Methods, Proteome completeness). Focusing on 62 KEGG modules (functional gene sets) present in at least half of the reference species we show that the *Achromatium* proteome recovered is at least 79 % complete. These modules include all main metabolic pathways involved in energy generation, nucleotide and amino acid synthesis, lipids biosynthesis, and specifically to *Achromatium,* sulfide oxidation (M00596) (Fig. S5). Notably, the experimental dataset lacks a module for ATP synthesis, F-type ATPase (M00157) entirely. However, it does contain the V-type ATPase which has been identified earlier in *Achromatium* genomes^3,6^, as well as two additional carbon fixation modules, the reductive pentose phosphate cycle (M00165) and the reductive citrate cycle (M00173), which were predicted to exist in our previous work^3,6^.

### Variant proteins and environmental cues and biophysics

For the 1456 OGs recovered in the MS-proteome, we extended the evaluation of variants described in the theoretical proteome analysis by a third category, the incubation temperature(s) in which the proteins have been found at least once. We will focus on the clusters established by biophysical predictions (Fig. 2, sequence cluster analysis in Fig. S6).

**Fig. 2:**
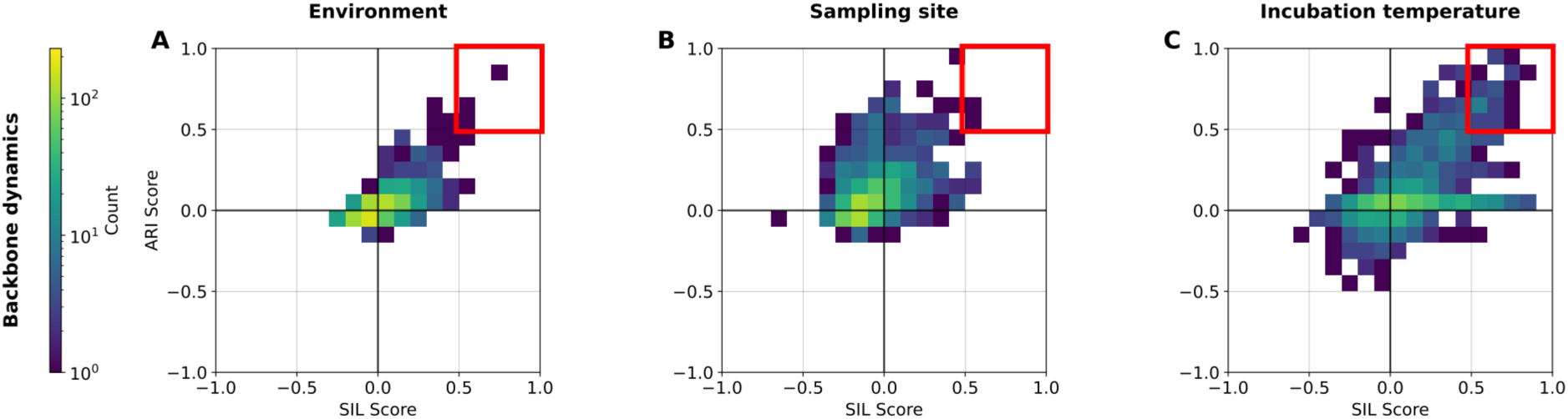
Variant assignment for expressed OGs is independent from environment (A) and sampling site (B) with trends preserved from Fig. 1. For incubation temperature (C), the distribution shifted towards positive values, with 54 OGs surpassing the 0.5 threshold (red box). Clustering based on difference of protein backbone dynamics. Scores for ARI and SIL range between -1 (worse than random), 0 (random) to 1 (perfect label matching/geometric separation). Number of OGs per bin is shown in color on a logarithmic scale.

For these filtered OGs, the trend established earlier remains, and no association between environment or sampling site and sequence-based clustering can be established with scores ranging between -0.2-0.1 for SIL and ±0.1 for ARI (Fig. 2A, B). Applying the 0.5 threshold for ARI and SIL, indicating geometric separation of the experimentally assigned clusters in agreement with the backbone-dynamics-derived clustering, identifies 54 OGs related to incubation temperature (Fig. 2C), with only single OGs in Fig. 2A and B surpassing the threshold. This increase over the environment and sampling location categories and illustrates the relevance of obtaining experimental proteome-level information for polyploid organisms.

Of the 54 OGs that were found to respond to incubation temperature, we identified 27 for which experimental evidence from other bacteria exists on protein-protein or protein-ligand interaction with information on the effect of mutations on these interactions (Tab. S 4). Below, we evaluated how the changes in predicted biophysical behavior connect to this further experimental information and the temperature-sensitive presence of variants in our work.

#### Show case 1: Disorder-mediated iron binding

OG0000334, representing the iron binding protein iscA, contains two distinct variants which differ strongest in protein disorder predictions in the region identified as crucial for the protein function using reference information.

iscA binds iron ions for iron-sulfur cluster formation and therefore contributes to a core metabolic process. In *Achromatium*, sequences from this OG were identified 76 times in the proteomic data across all conditions and timepoints. These sequences were mapped to two clusters established in the theoretical proteome (Fig. 3A-C), one of which was not seen in the high-temperature condition while the other was seen in all conditions (Fig. 3E-G). Most

**Fig. 3:**
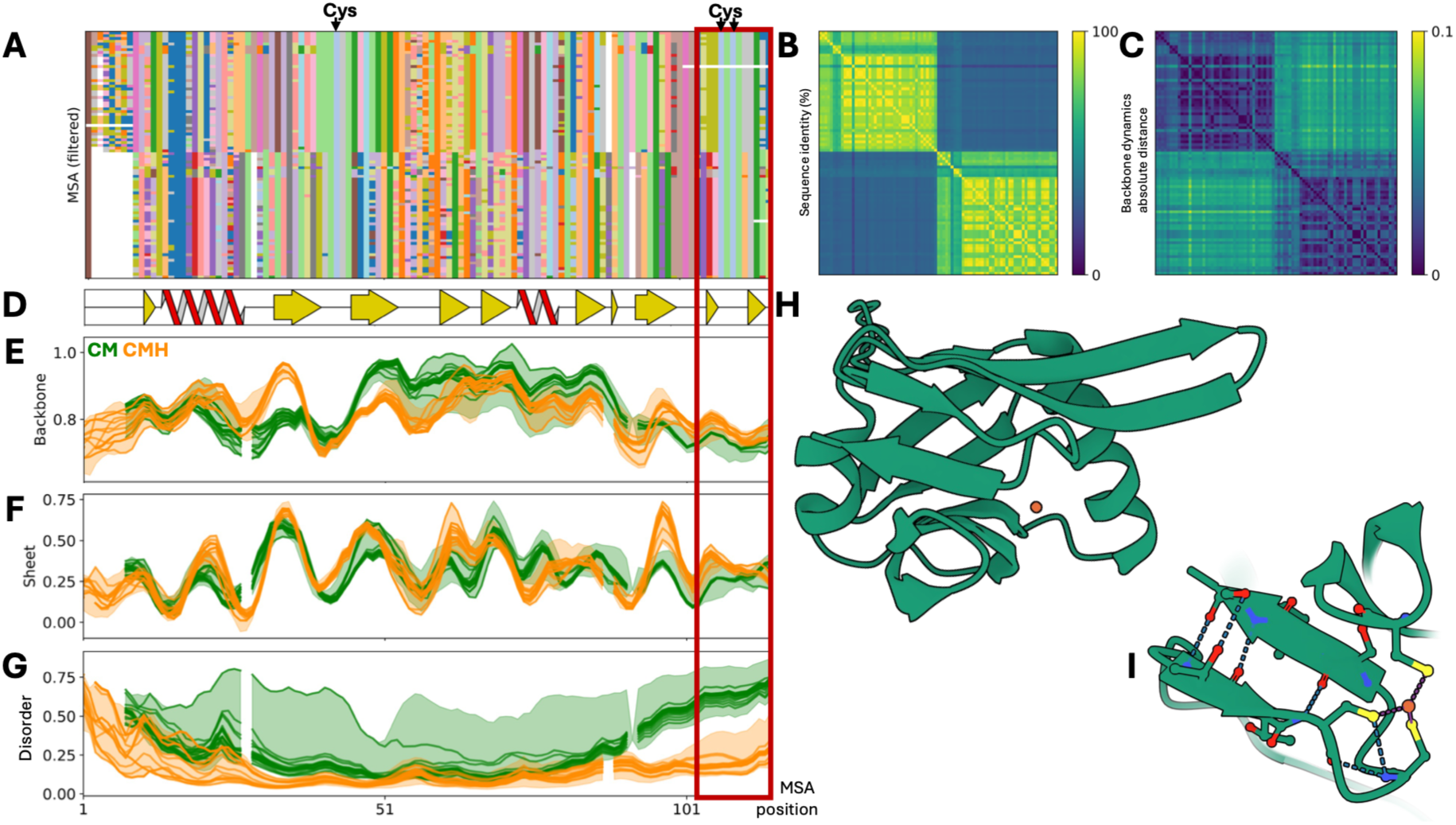
Iron-binding protein iscA variants differ in disorder and beta-sheet propensity. (A) Filtered MSA of two clusters with presence in experimental data amended with reference sequence of P0AAC8 and conserved functionally relevant cysteines indicated above. (B) Pairwise sequence identity matrix ordered to match MSA highlighting within versus between cluster sequence identity. (C) Distance matrix calculated with the absolute difference per position for backbone dynamics predictions normalized by alignment length. (D) Consensus secondary structure elements a-helix (red) and ß-sheet (yellow) from predicted protein structures mapped onto MSA. (E-G) Predictions of biophysical propensities mapped onto the MSA. Predictions for sequences identified in experimental data are shown as lines, the shaded area indicated the range of predictions for the members of the cluster. Clusters/Sequences identified in all conditions (CMH) in orange, with CM conditions in green. (E) Prediction values for backbone dynamics. Values below 0.8 indicate a flexible region, above indicate rigidity. CM variant is more rigid between positions 45-90, but more flexible in the terminal regions than CMH. (F) Prediction values for ß-sheet. Higher values indicate that the region is more likely present in this secondary structure. The CMH variant shows increased values from position 95 on. (G) Prediction values for disorder. Regions above a threshold of 0.5 are considered disordered, such as the CM C-terminal region. (H) AlphaFold3 prediction of CMH-variant protein sequence (green) co-folded with an iron cation (orange). (I) Focused view of (H) showing iron cation binding with the three conserved cysteines and ß-sheets stabilizing the cysteines relative position towards each other. This region is also indicated in the MSA (A, D-G) view in a red box.

*Achromatium* genome assemblies contained both protein variant sequences. P0AAC8, iscA protein in *E. coli*, was identified as a reference in the UniProt database. Cysteines at positions 99 and 101 are crucial to the protein function, while a mutation of cysteine 35 leads to a decrease in iron-binding activity^26^. These positions are also strictly conserved in the *Achromatium* protein (Fig. 3A). Modeling the ligand binding in 3D using AlphaFold3 (Fig. 3H) revealed that all three cysteines are located in the same area and bind the iron ion together, despite the distance in sequence-view (Fig. 3I). The modelling also reveals ß-sheets on both ends of the loop containing the two crucial cysteines, which holds them close proximity (Fig. 3H). In the CMH variant (orange color), this region is more rigid (Fig. 3E) and more likely to form a ß-sheet (Fig. 3F). In the CM variant (green), however, this region is predicted to be disordered (Fig. 3G), making it overall less likely that these residues appear in this conformation.

#### Show case 2: Protein recognition and rigidity

OG0001449 represents the ribosomal protein rpsI, which in the case of *Achromatium* harbors three variants which differ in rigidity in the region responsible for protein-protein recognition.

The rpsI protein, a central member of the small ribosomal subunit (uS9), was identified 37 times in the proteomic data across all conditions and timepoints. These sequences mapped to three clusters established in the theoretical proteome (Fig. 4A-C), one of which was not seen in the high-temperature condition H with the other two being expressed in all conditions (Fig. 4E,F). Assemblies have one or two variants of this protein, however, not all variants were in the experimental dataset. P0A7X3, rpsI protein in *E. coli*, was identified as a reference in the UniProt database. The deletion of C-terminal region leads to reduced growth of *E. coli* at low temperatures^27^. We also identified an experimental structure of the complete 70S ribosome (PDB ID 9SYH^28^), which allowed us to understand how rpsI interacts with proteins and RNA in the complex. The sequence of P0A7X3 (identical to 9SYH) was added to the filtered MSA to transfer these regions of interest, which are indicated above the MSA (Fig. 4A). The disordered region (Fig. 4F, second red box) is shown to extend through the ribosomal RNA (Fig. 4G). In the CMH variants, proteins are predicted to be more disordered in a larger region than the CM variants. Other residues interacting with the rRNA are located in the second, third and fifth ß-sheet (Fig. 4D). Further, the predicted backbone rigidity increased in CM variants in the region interacting with rpsH (Fig. 4E, first red box). Considering the 3D structure (Fig. 4H), we see the strictly conserved arginine in position 41 make connection with rpsH. The unstructured tail of rpsI is close to the helix indicated in (Fig. 4E, first red box).

**Fig. 4:**
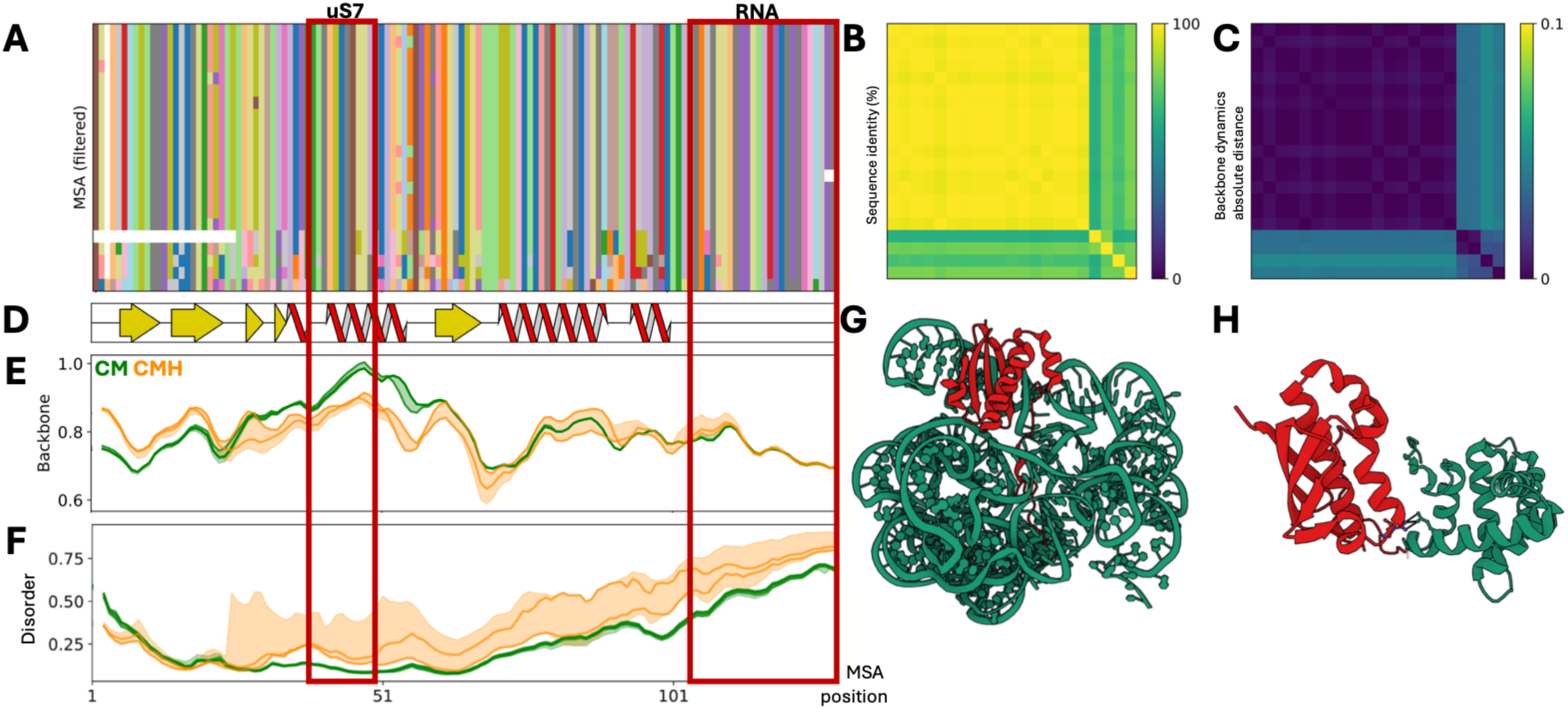
Small ribosomal subunit protein rpsI differs in disorder and backbone dynamics. (A) Filtered MSA of clusters with presence in experimental data amended with reference sequence of P0A7X3 and regions identified for protein interaction indicated above in a red box. (B) Pairwise sequence identity matrix ordered to match MSA highlighting within versus between cluster sequence identity. (C) Distance matrix calculated with the absolute difference per position for backbone dynamics predictions normalized by alignment length. (D) Consensus secondary structure elements a-helix (red) and ß-sheet (yellow) from predicted protein structures mapped onto MSA. (E,F) Predictions of biophysical propensities mapped onto the MSA. Predictions for sequences identified in experimental data are shown as lines, the shaded area indicated the range of predictions for the members of the cluster. Clusters/Sequences identified in all conditions (CMH) in orange, with CM conditions in green. (E) Prediction values for backbone dynamics. Values below 0.8 indicate a flexible region, above indicate rigidity. CM variant is more rigid between positions 30-60. (F) Prediction values for disorder. Regions above a threshold of 0.5 are considered disordered, such as the C-terminal region. CMH disordered region may start at position 90, whereas CM disordered region only starts at position 105. (G,H) Excerpt from experimental structure of E. coli 70S ribosome (9SYH). (G) rpsI (red) interacting with 16S rRNA (green) via the disordered tail (red box in the MSA view (A,D-F)) and ß-sheets. (H) rpsI (red) helical region (red box in the MSA view (A,D-F) interacting with rpsH, also known as uS7 (green).

#### Show case 3: Protein recognition

OG0001414 is representing the stringent starvation protein sspB. This adaptor protein is delivering the rseA factor to clpX protease for degradation, thus triggering the cell envelope stress response^29^. *Achromatium* harbor two distinct variants which differ in secondary structure and protein disorder predictions in the regions identified for protein-protein recognition.

The sspB protein was identified 9 times in the proteomic data across cold and medium conditions and all timepoints. These sequences mapped to two clusters established in the theoretical proteome (Fig. 5A-C), one of which seen only in the low-temperature condition C and the other was seen in CM conditions (Fig. 5 E-G). P0AFZ3, sspB protein in *E. coli* was identified as a reference in the UniProt database. Mutations in positions 52, 59 and 74 lead to a decrease of rseA binding^30^. Additionally, the C-terminal region, from position 128, was indicated as disordered and for binding of protease clpX in the UniProt entry. The experimental structure of the sspB dimer bound to rseA (PDB ID 1YFN^30^), allowed us to understand how sspB interacts with itself in dimerization and rseA. The sequence of P0AFZ3 (identical to 1YFN) was added to the filtered MSA to transfer the locations of these regions indicated above the MSA (Fig. 5A). The first red box highlights the first a-helix (Fig. 5D), position 10-23, through which two sspB proteins interact to form a dimer (Fig. 5H). In the CM variant, the helix prediction value drops for the last helix turn (4 residues) sheets (Fig. 5E). The tryptophan in position 20, as well as the aspartic acid in position 23, remained conserved and form hydrogen bonds with the other sspB protein. The second region of interest (Fig. 5A, position 50-76) contains the three positions responsible for rseA binding, framed by ß-sheets (Fig. 5D). The increase for a-helix match the decrease for ß-sheet prediction values especially for region 60-65 (Fig. 5E, F) in the C variant. In the 3D structure (Fig. 5I) the loop between second and third ß-sheet, which contains position 59, is shown as a helix turn. Finally, the C-terminal region in the reference sequence is much longer, causing the gaps in alignment and biophysical predictions. The region is predicted to be disordered in the CM variant, but the C variant (Fig. 5G). In the experimental 3D structure, this region could not be resolved. Therefore, we predicted the 3D structure using ESMfold (Fig. 5J), which shows most of the region unstructured with a short helix at the start.

**Fig. 5:**
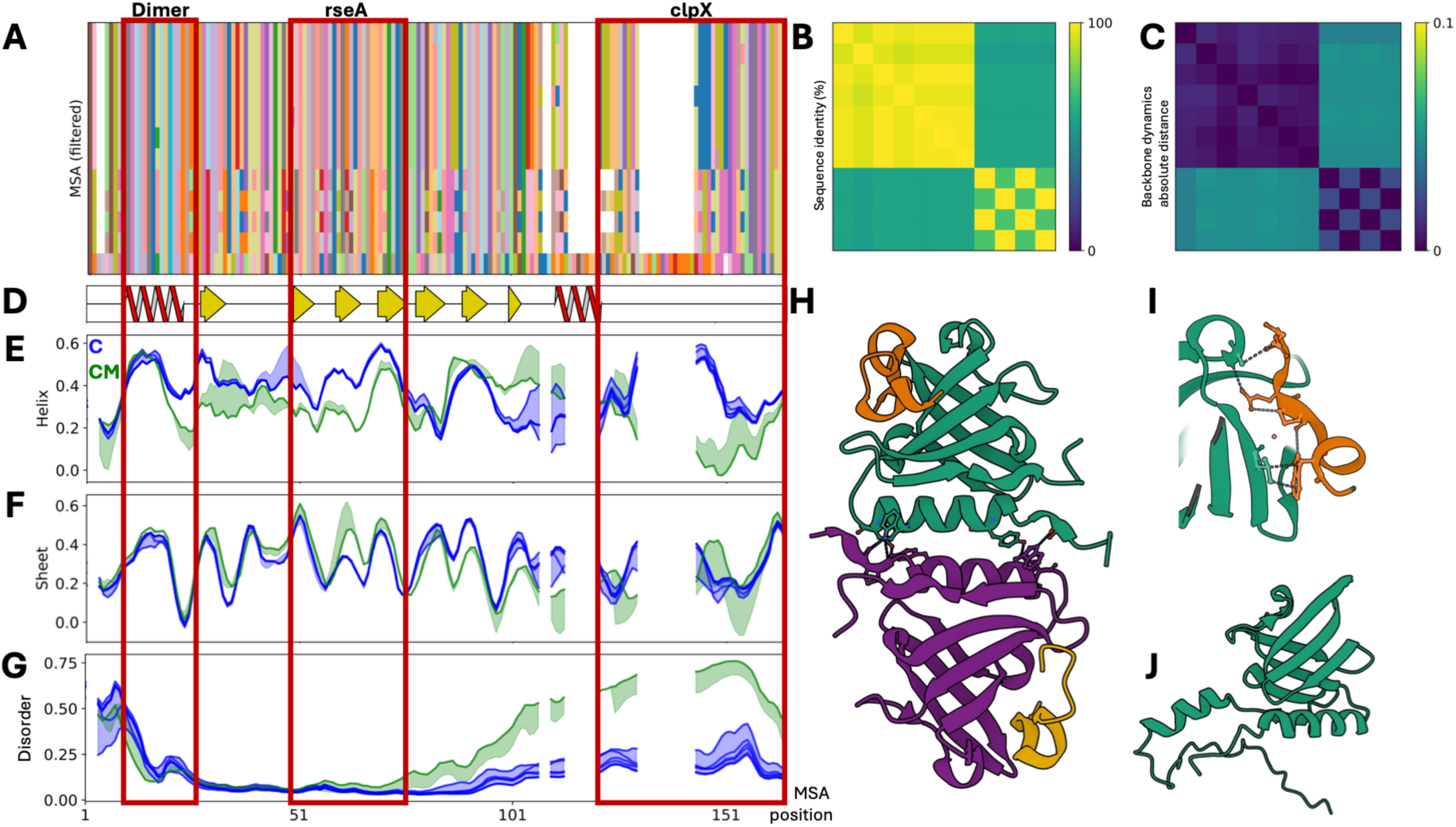
Starvation protein sspB variants differ in disorder and secondary structure predictions. (A) Filtered MSA of two clusters with presence in experimental data amended with reference sequence of P0AFZ3 and regions identified for protein interaction indicated above in a red box. (B) Pairwise sequence identity matrix ordered to match MSA highlighting within versus between cluster sequence identity. (C) Distance matrix calculated with the absolute difference per position for backbone dynamics predictions normalized by alignment length. (D) Consensus secondary structure elements a-helix (red) and ß-sheet (yellow) from predicted protein structures mapped onto MSA. (E-G) Predictions of biophysical propensities mapped onto the MSA. Predictions for sequences identified in experimental data are shown as lines, the shaded area indicated the range of predictions for the members of the cluster. Clusters/Sequences identified only in cold conditions (C) in blue, with CM conditions in green. (E, F) Prediction values for secondary structure propensities mapped onto the MSA. Higher values indicate that the region is more likely present in this secondary structure. (E) Prediction values for a-helix. C variant predictions are higher overall. (F) Prediction values for ß-sheet. The CM variant shows increased values for the third ß-sheet at position 60-65. (G) Prediction values for disorder. Regions above a threshold of 0.5 are considered disordered, such as the CM C-terminal region, whereas for the C variant no disorder is predicted. (H) Experimental structure of E. coli sspB dimer (green, violet) bound to rseA (1YFN) showing proteins interaction via the first helix. (I) Focused view of (H) showing rseA (orange) and sspB (green) interacting at the second ß-sheet and turn. (J) Prediction of the full 3D structure with ESMfold.

## Discussion

*Achromatium* is a genus of giant bacteria with an unusual amount of consistently maintained chromosomal diversity^1^. In this study, we illustrate how essential proteomics is to investigate whether this genomic diversity serves to improve and expand *Achromatium’s* response to environmental signals in the functional realm of proteins. By coupling sequence identity and biophysical metrics we showed that sequence divergence within OGs is not random. Instead, many groups contained distinct clusters of sequence variants, consistent with the maintenance of different protein forms rather than the unrestricted accumulation of substitutions across chromosomes. Despite the extreme diversity found in the *Achromatium* proteome, residues with functional importance are often strictly conserved across the whole OG. By calculating the proteins biophysical properties from individual sequences, we were able to detect shifts in the local sequence context, such as mutations in the surrounding residues that can modulate electrostatic interactions of the protein with itself, ligands or other proteins.

We demonstrated how our analysis can extend beyond variant detection to generate well-founded hypothesis on the mechanisms that make variants specific to an environment. The three showcase proteins (iscA, rpsI, sspB) illustrate how temperature-dependent variant expression operates in *Achromatium*. Protein variants maintain their core functions while their biophysical properties adjust, e.g. through a change in protein backbone flexibility^21,31^. In the iron binding protein (iscA) the variant expressed also at high temperatures has an increased likelihood of forming a second ß-sheet at the C-terminus of the protein. This ß-hairpin structural motif has been shown in other enzymes to coordinate functional residues, such as the two cysteines responsible for binding the iron ion^32,33^. The increase of energy in the system due to the elevated temperature may cause the effect of the increase in predicted disorder in variant expressed at lower temperatures to compound, making this ideal conformation less likely, and therefore the variant less suitable^34,35^. Proteins directly identified in the MS proteomics served as anchors to extend experimental annotations to all members of the variant cluster within the OG, thus allowing us to encompass more of *Achromatium*’s diversity in our analysis. This inference strategy was especially important for the analysis of less detected proteins, such as rpsI and sspB. The increase in intrinsic disorder, which we observe in the generalist variant of the small ribosomal protein (rpsI) may enhance binding promiscuity with the many biomolecules involved in the ribosome during translation^27,36^. Finally, the efficiency of sspB substrate delivery to the clpX protease depends on appropriate spacing between them^37^. The increase in helix propensity and decrease in disorder we observe in the cold variant allows sspB to maintain distance from clpX and prevent steric clashes.

We identified temperature-associated variants in approximately 5 % of the OGs detected in the proteome. The strongest patterns of selective variant detection were observed in the 30 °C samples, despite the relatively small temperature difference between M and H conditions. As Lake Stechlin, from where the cells originate, rarely exceeds 25 °C^38^, this may be a tipping point in triggering the expression of heat-relevant variants. This threshold-based mechanism is consistent with heat shock systems in other organisms, which exhibit bistable switching rather than gradual induction for a small part of their proteome^39^. The expression of temperature-responsive protein variants we show here is consistent with the general principle of thermal adaptation. Most protein variants were detected in both the cold and room-temperature (23 °C) conditions. The remaining 95 % OGs may still include variants responding to other environmental factors such as salinity, or temperature sensitive proteins that were not sufficiently sampled in the current dataset. Indeed, fewer proteins and cells were recovered from the 30 °C treatment, making comparisons involving this condition more sensitive to sampling effects.

The limited representation of *Achromatium* in public databases restricts peptide identification and makes an organism-specific sequence database essential for comprehensive proteome analysis. Only by integrating the non-redundant version of the *Achromatium* atlas into the workflow were we able to significantly increase the proteome coverage, achieving a near 20-fold increase compared to the standard workflow. The high sequence similarity among related protein variants presents an additional analytical challenge, as many peptides may be shared among several sequences and thus cannot be uniquely assigned. Conversely, sequence divergence can generate unique peptides that distinguish individual variants, providing evidence for their condition-specific expression. Interpreting the data at both the individual protein-sequence and ortholog-group levels therefore enables variant-specific evidence to be retained while also accounting for closely related proteins that cannot be unambiguously distinguished. Nevertheless, the absence of a detected variant does not necessarily indicate its biological absence, and analysis of pooled cells cannot establish whether different variants originated from distinct chromosome copies within the same cell or from different cells within the sampled population. Advances in meta-proteomics may in the future provide innovative ways to address the challenges outlined here^40^. In particular, continued improvements in the sensitivity of mass spectrometry-based proteomics may enable in-depth proteome characterization at the single cell level^41^. In the absence of a validated reference genome or proteome, conventional assessments of proteome completeness were not applicable. However, with a coverage of 64 % of the OGs in the *Achromatium* atlas and a coverage of 79 % of the KEGG metabolic pathways broadly conserved in the *Chromatiaceae* family, our experimental data provides a qualitative benchmark for future *Achromatium* proteomes.

We show that variant presence relates to transient changes in water temperature, suggesting that this feature likely extends to other varying environmental signals. In contrast, the diversity of the *Achromatium* cannot be connected to the broader, more stable, environmental conditions like the sampling environment, i.e. marine or freshwater. This is in agreement with previous comparisons of the functional repertoires of marine and freshwater *Achromatium* genomes which did not identify environment specific functions^3,6^. Similarly, these variants are not associated with sampling location indicating that shared ancestry is not the driving force in the sequence diversity in *Achromatium*.

Our results provide evidence that the hyper-polyploid species *Achromatium* maintains distinct protein-sequence variants which are expressed selectively as a fine-tuned response to transient environmental cues such as temperature. This opens the door to speculate whether *Achromatium* is the only prokaryote with this regulatory mechanism, when it has been shown that other unicellular giant bacteria also contain significant genomic diversity^5^.

## Methods

### Selective incubation

Fresh *Achromatium* cells were collected in May 2025 in lake Stechlin near Neuglobsow, Brandenburg, Germany (53° 9′ 5.59″ N; 13° 1′ 34.22″ E) as previously described^1^. Samples were incubated at three temperature conditions: at 7 °C (cold - C), at room temperature, measured on average at 23 °C +/- 3 °C (medium - M), and 30 °C (hot - H). All samples were kept in the dark to prevent algae growth. Cells were collected after 1 day (T0, only at M), after 2 weeks (T1) and after 4 weeks (T2). Each condition was incubated in triplicate. The cells were collected using the previously established protocol^1^. Cells were transferred into 1.5 mL Eppendorf tubes and gently spun down with a tabletop centrifuge. Supernatant was discarded. Cells were resuspended in 1 mL sterile filtrated lake water and frozen at –20 °C. Finally, the cells were lyophilized under vacuum.

### Protein amount estimation and sample preparation

Cell pellets were lysed using Laemmli buffer containing 10 mM Tris-HCl (pH 6.8), 10 % glycerol, and 5 % SDS. Cell lysis was further facilitated using a Bioruptor sonication system (Diagenode) for 10 min at 4 °C, with alternating cycles of 30 s sonication (ON) and 30 s rest (OFF). Protein concentrations in the resulting lysates were determined using the Pierce 660 nm Protein Assay Kit (Thermo Scientific), according to the manufacturer’s instructions. High-purity bovine serum albumin (BSA) was used to generate the standard calibration curve.

The lysates were subsequently processed for bottom-up proteomic analysis using a stacking-gel sample preparation^42^. Based on the measured protein content (Tab. S 3), the samples were divided into two groups. For samples containing more than 10 µg of protein in the volume available for gel loading, 10 µg of protein were loaded. For samples containing less than 10 µg of protein, 2 µg was loaded. Stacked proteins were reduced and alkylated before Trypsin/Lys-C digestion using an enzyme-to-protein ratio of 1:25 for 15 h at 37 °C under agitation at 600 rpm. The resulting peptides were extracted from the gel pieces and stored at −80 °C until further analysis.

### LC-MS/MS analysis

The frozen peptide samples were vacuum-dried and subsequently resuspended in 2 % acetonitrile (ACN) containing 0.1 % formic acid (FA) to a final concentration of 100 ng/µL. A pooled sample was prepared by mixing 5 µL of each resuspended sample. This pooled sample was used exclusively to design the acquisition method and was not included in the subsequent quantitative analysis.

Peptide separations were performed using a nanoElute liquid chromatography system equipped with a 25 cm IonOpticks analytical column. For method design, the pooled sample was separated using a 100 min linear LC gradient ranging from 2 to 80 % B (ACN/0.1 % FA) at a flow rate of 0.2 µL/min. For the final quantitative analysis, 200 ng of peptides from each individual sample were injected and separated using an 80 min linear LC gradient at the same flow rate.

MS analyses were performed using a timsTOF Pro 2 mass spectrometer (Bruker Daltonics). The pooled sample was initially analyzed in DDA-PASEF^43^ mode to characterize the precursor-ion distribution across the mass-to-charge and ion-mobility dimensions. The ion-mobility range was set to 0.70–1.40 V·s·cm⁻², and MS spectra were acquired over a m/z range of 100–1700. The resulting precursor-ion distribution was used to define the ion-mobility range and optimize the isolation-window layout of a sample-specific narrowPASEF^44^ method, a tailored variant of diaPASEF^45^. The optimized narrowPASEF method was subsequently applied to the individual samples for quantitative analysis. The acquisition scheme comprised 26 MS/MS isolation windows, each 25 Da wide, covering a m/z range of 400.4–1050.4 and an ion-mobility range of 0.71–1.29 V·s·cm⁻². The accumulation and ramp times were both set to 100 ms. Collision energy was linearly increased from 20 eV at 0.60 V·s·cm⁻² to 59 eV at 1.60 V·s·cm⁻². Each acquisition cycle lasted 1.27 s and consisted of one full MS1 scan followed by 26 MS/MS windows.

### Computational proteome analysis

Below we describe the data analysis scheme used to assess and interpret the diversity of ortholog proteins in *Achromatium*. The numbered steps in Fig. 6 are referred to in the text in square brackets.

**Fig. 6:**
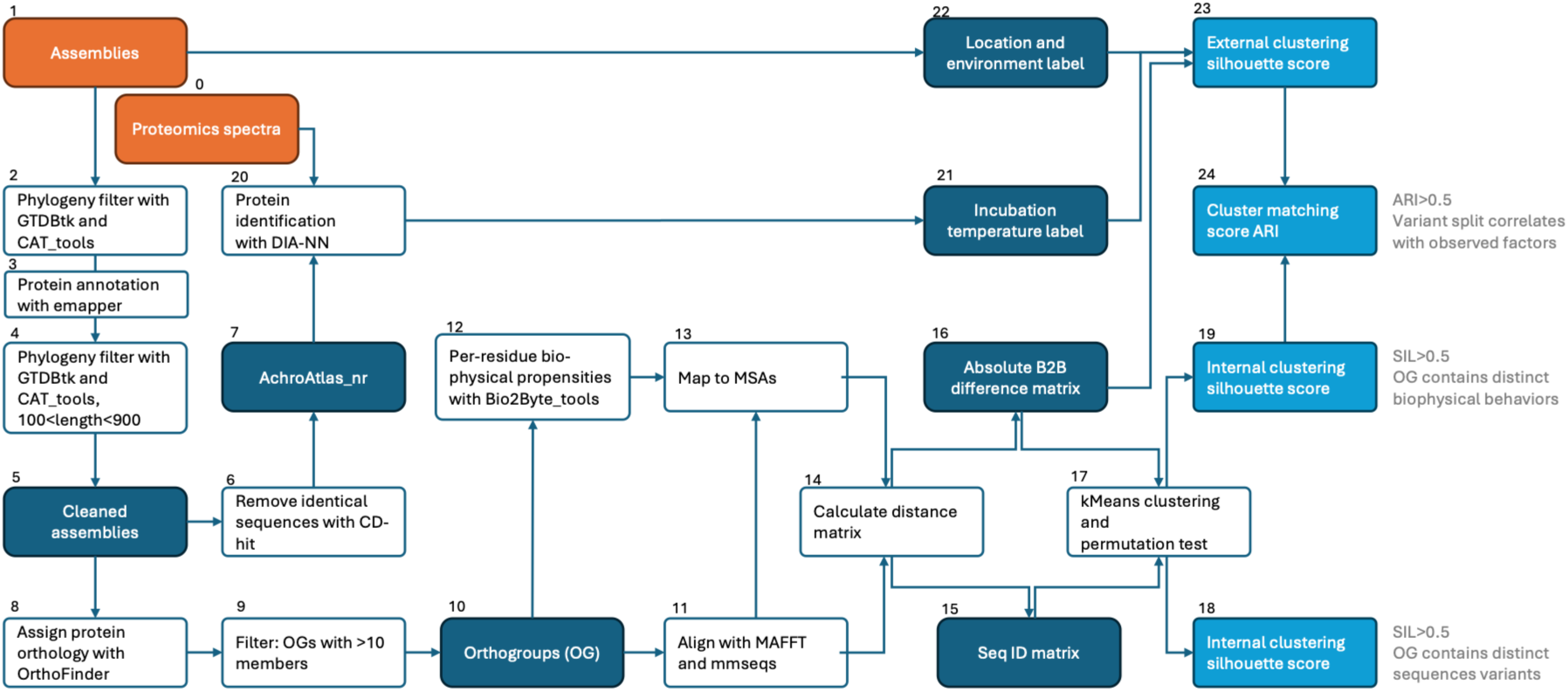
Numbered analysis scheme combining genomic and MS-proteomic data to identify OGs with temperature-sensitive variants.

#### Data collection and cleaning

[1] 13 publicly available genomes^7,22,23^, taxonomically identified in NCBI as *Achromatium* from various environments, were added to 10 metagenomes^6^ and 32 single cell assemblies^1,6^ from Lake Stechlin samples. Additionally, we integrated 2 assemblies from samples from mangroves at Manche-à-Eau lagoon, Guadeloupe (Rémy et al. in preparation). [2] The phylogenetic analysis of all assemblies was carried out using GTDBtk v2.4.1^46^ with database release 226 (April 2025). 3 genomes from NCBI were annotated as Alphaproteobacteria and 1 genome identified as *Thermochromatium* were excluded from further analysis. The contigs of the remaining assemblies were assessed for contamination using the Contig Annotation Tool CAT v6.0.1^47^, resulting in the exclusion of on average 10 % contigs per assembly. [3] The cleaned genomes were annotated using eggnog mapper^48^ (emapper v2.1.9) and database 5.0.2. For 2 assemblies, no proteins were called. [4] Finally, a sequence length filter of >100 and <900 residues was applied to remove fragmented protein sequences. [5] This finally resulted in a dataset of 51 assemblies (Tab. S 1) containing ∼180K sequences. [6] Using CD-Hit^49^, [7] an *Achromatium* atlas was established with 150K unique sequences in FASTA format. These sequences are considered the theoretical proteome of *Achromatium*. Two assemblies containing each less than 100 proteins were excluded from further analysis. For the assessment of protein orthology, [8] OrthoFinder^17^ was used with standard settings with a custom phylogenetic tree [9] Only OGs with more than 10 members were considered for further analysis, which resulted in 2259 OGs [10].

#### Protein identification and quantification

[0] The raw data were processed using DIA-NN version 2.2.0^50^. A predicted spectral library was first generated from the *Achromatium* atlas [7]. This library was amended to the standard UniProt library and was subsequently used for peptide and protein identification and quantification from the experimental datasets. The number of allowed missed cleavages was set to 1. Carbamidomethylation of cysteine residues, Methionine oxidation (Ox(M)) and N-terminus acetylation (Ac(N-term)) were permitted as modifications. Mass accuracy was set to 1.5 × 10⁻⁵ for MS2 and 5 × 10⁻⁶ for MS1, and a scan window of 10. The false discovery rate (FDR) was set to 1 % at precursor level, and match-between-runs (MBR) was enabled. [20] The protein group matrix generated by DIA-NN, with a global protein group FDR threshold of < 1 %, was used for the downstream data analysis.

#### Distance matrices and clustering

This part of the analysis is implemented in a Nextflow^51^ pipeline available on GitHub https://github.com/slheidig/proteinfam_pca_nf. It uses Docker and Apptainer for dependency management and is optimized to use HPC infrastructure to parallelize calculations.

[11] We considered two types of sequence alignments to calculate the sequence distance matrix: MAFFT^52^, which produces multiple sequence alignments, and MMseqs2^53^, which produces pairwise sequence alignments and directly provides sequence identities. For MAFFT alignments, [14] sequence identity is calculated for each pair treating positions where both sequences have a gap, positions where only 1 sequence has a gap as a mismatch and comparing all other positions. The final score is normalized by the number of valid positions. This results in a sequence distance matrix [15].
[12] For each sequence, 7 biophysical propensities are calculated using the b2bTools^16^. However, in following only the feature containing the dynamics of the protein backbone are considered. To compare these per-residue features between sequences of the same OG, [13] the predictions are mapped onto the sequence alignments. The biophysical distance matrix [16] is calculated similarly to the sequence matrix by calculating the absolute difference between each residue and normalizing by the number of valid positions.
[17] To each distance matrix we apply the k-means algorithm implemented in the sklearn python package^54^. We test between k=2 until k=10. The clustering is evaluated using the silhouette score (SIL), a measure of the geometric separation of all points within versus outside a cluster. Using permutations, a p-value is assigned to this reference free score [18,19]. SIL ranges from -1 to 1, with a score of 0 denoting randomness and 1 perfect separation.

To interpret these clusters, external information was integrated from two sources. Information on the sampling location (i.e. Lake Stechlin) and environment at the location (e.g. Freshwater) was assigned at assembly level [22]. Therefore, every sequence has such a label. Proteins that have been identified in the experimental proteomics data [0,20], which is approximately 10 % of all proteins in the theoretical proteome in 1500 OGs, can be [21] assigned the incubation temperature label, which denotes if the protein was found in the cold (C), room temperature (M) or hot (H) condition, or any combination between them. These external labels are assigned to the distance matrix [16] and evaluated using the SIL. [23] Finally, the overlap between the external and internal clustering is evaluated using the adjusted Rand index (ARI). The ARI also scales between -1 and 1, with 0 denoting random assignments and 1 perfect match between both labeling strategies.

#### Cluster selection

While both matrices are calculated on the basis of multiple sequence alignments, they capture slightly different effects. [18] The sequence identity score captures if the OG contains distinct protein variants. [24] The biophysical score captures if those variants have the potential for different behavior. The cutoff for both is SIL>0.5. Additionally, the OGs are filtered by an ARI >0.5 for the incubation temperature category. This results in 54 OGs (Tab. S 4).

#### Proteome completeness

The sequences of all proteins identified in the experimental proteome were mapped back to the *Achromatium* atlas, showing experimental evidence for 1456 of the 2259 OGs (64 % coverage). To assess the coverage of the standard metabolic pathways, the experimental proteome was submitted to GhostKOALA^55^ to map them to KEGG identifiers (KID). 53 % of sequences were mapped to a KEGG ID (KID). Using the KEEG website, the pathways these KID belonged to were recovered as modules (metabolic gene sets). KIDs were assigned to 195 modules. A module completeness (KID found/KID expected) threshold of 50 % was established, reducing the list to 119 modules. The KEGG database in July 2026 contained 52 datasets of *Chromatiaceae* family, which *Achromatium* belongs to. This reference set contains 143 modules, with 84 modules found in both the reference and experimental datasets (58 % coverage). Considering only the modules present in more than half the *Chromatiaceae*, the reference set was reduced to 62 modules (79 % coverage). We also prepared reference set of modules expected to be complete in *Bacteria*. This general reference set contained 408 modules (28 % coverage), with only 49 common modules (84 % coverage).

### Reference published variant effects

Functional annotations for each of the 54 selected OG are established from the consensus of emapper results and verified by running BLAST through the Uniprot portal^56^ on the longest sequence per cluster that were found in experimental data. The consensus protein name is used to search well annotated orthologs on Uniprot with mutagenesis results or experimentally validated binding sites or protein-protein interaction interfaces. Structures are predicted using ESMfold^57^ for additional verification of the match between reference protein and the *Achromatium* data. The reference sequence is aligned to the MSA of the OG to identify functionally relevant positions in the OGs. The MSA is filtered for variant clusters with presence in the proteomics dataset, as well as the prediction of biophysical behaviors such as backbone dynamics, sheet propensity or disorder. We map secondary structure elements onto the MSA using DSSP^58^. This information is plotted per variant per residue using the MSA as a shared reference frame. Finally, we consider the relative position of the residues in 3D space using the structure predictions. We also consider experimental structure information using the PDB^59^. Where no direct experimental evidence is available, we use AlphaFill^60^ and AlphaFold3^61^ to extend information of ligand binding and protein-protein interfaces. Visualization and analysis of protein structures is carried out in Mol*Viewer^62^. Combining this spatial analysis with the per residue predictions from the b2bTools allows to present well founded hypothesis on the mechanisms that make variants specific for environmental conditions.

## Data/ Code availability

Data and analysis scripts for the computational proteome analysis are available on Zenodo (10.5281/zenodo.22645072). The clustering analysis code is available on GitHub (https://github.com/slheidig/proteinfam_pca_nf). The mass spectrometry proteomics data have been deposited to the ProteomeXchange Consortium via the PRIDE^63^ partner repository with the dataset identifier PXD083615.

## Acknowledgments

The authors wish to thank the department of Plankton and Microbial Ecology at the Leibniz Institute of Freshwater Ecology and Inland Fisheries (IGB) for allowing access and to Lake Stechlin for sample collections. The authors are grateful to Prof. J.F. Flot (Ecological & evolutionary genomics lab, ULB) for his valuable discussions and guidance. The computational resources used in this work were provided in part by the VSC (Flemish Supercomputer Center), funded by the Research Foundation—Flanders (FWO) and the Flemish Government.

## Author contributions

S.L.H. and D.I. conceived the study. S.L.H. and D.I. designed the experiments and performed the selective incubation experiment. D.I., J.M.V., O.G. and P.R. sequenced and assembled *Achromatium* genomic data. S.L.H. curated the *Achromatium* protein atlas. S.L., J.B.R. and C.C. performed MS-proteomics and data interpretation. S.L.H., D.I. and W.V. analyzed and interpreted the data. S.L.H. wrote the manuscript with input from all authors. D.I. supervised the study and secured funding.

## Ethics declaration

The authors declare no competing interests.

## Supplementary materials

**Fig. S1:**
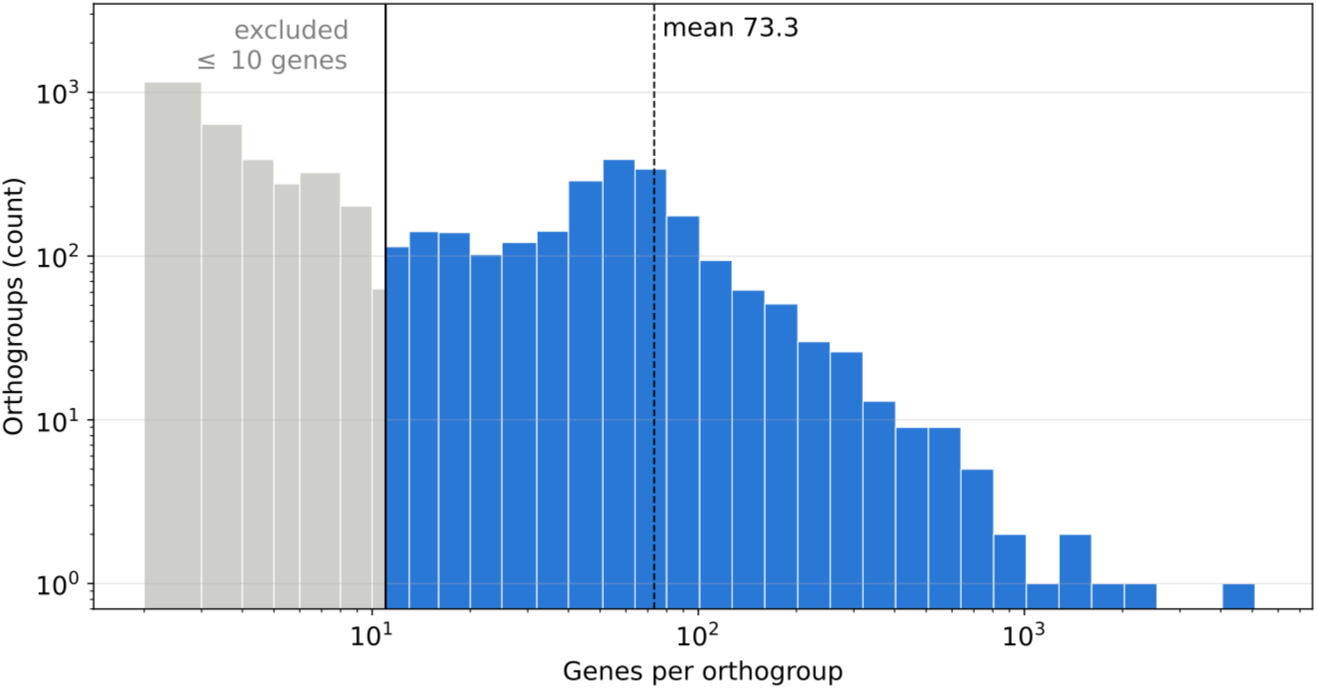
Distribution of gene counts per group of ortholog proteins. OGs with 10 members or less were excluded from the analysis. The largest bin is 51-64 members with 389 OGs. OGs contain on average 73.3 sequences, with 6 OGs containing more than 1000 sequences.

**Fig. S2:**
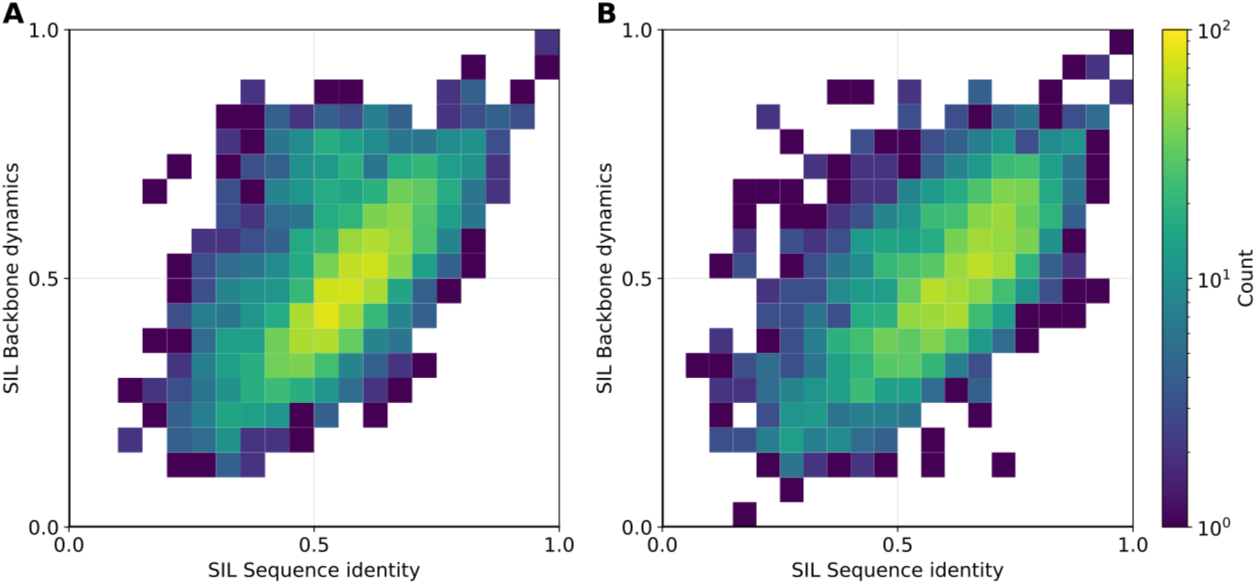
Distribution of SIL scores for all OGs for k-Means clustering based on sequence identity or difference in backbone dynamics. Both (A) MAFFT, a global alignment method and (B) MMSEQS2, which performs pairwise alignments support the existence of distinct variants in most OGs at a similar level. The threshold of 0.5 is surpassed by 981 and 1,031 OGs using MAFFT or MMSEQS2 respectively.

**Fig. S3:**
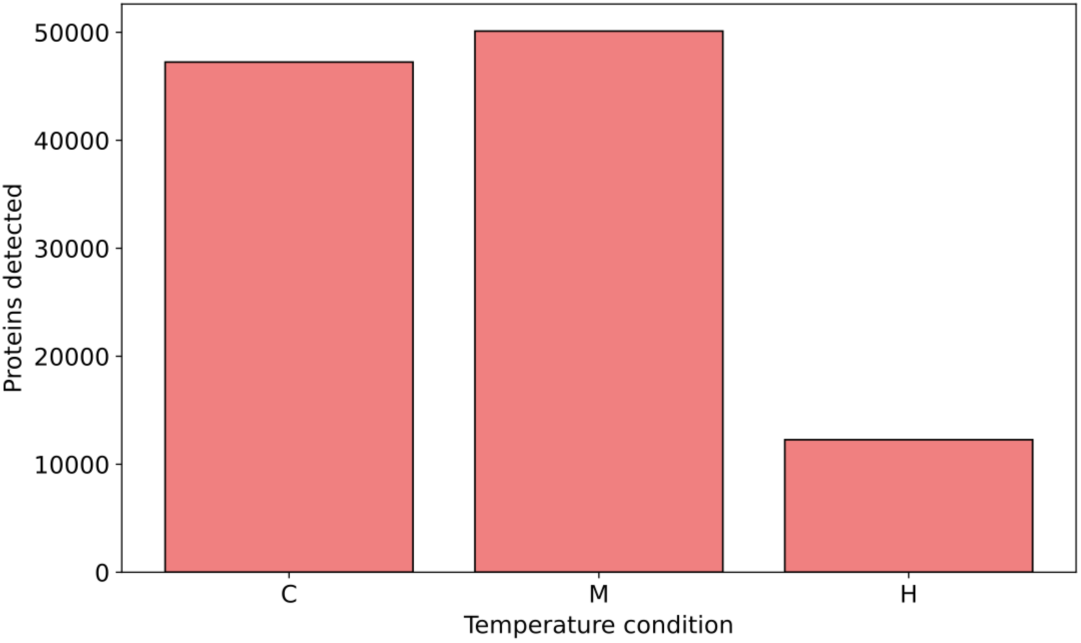
Distribution of unique proteins identified in each temperature condition. Cold [C], medium [M] and hot [H] denote incubation temperatures. One protein can occur in more than once, i.e. it is detected in both in C and M condition and replicates.

**Fig. S4:**
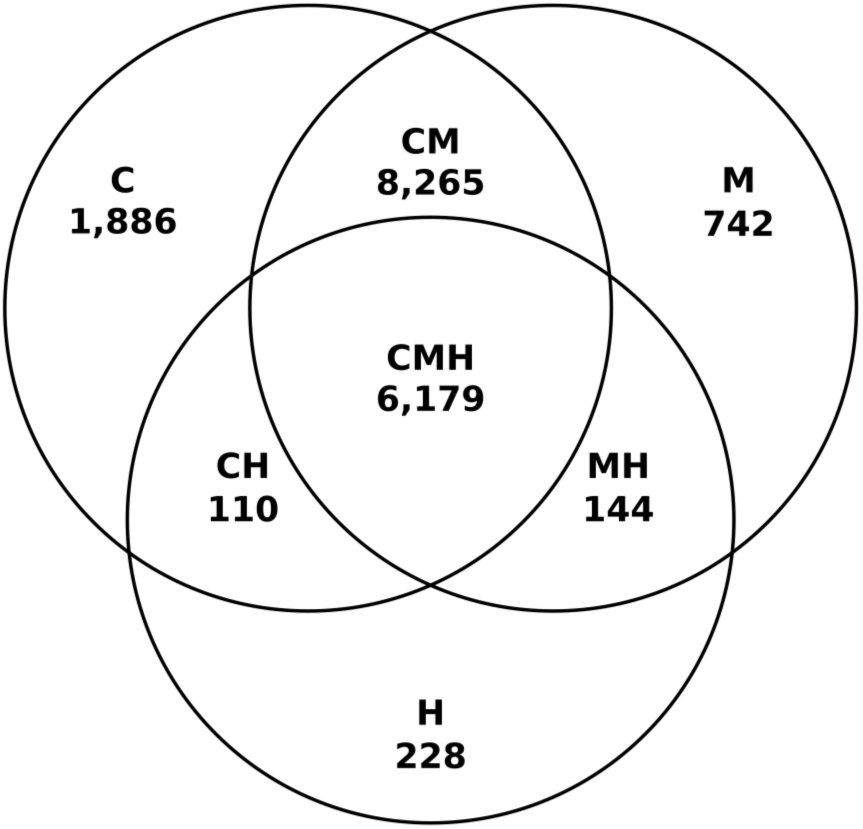
Venn diagram of distribution of unique proteins detected per temperature category. Proteins occur only once. Cold [C], medium [M] and hot [H] denote incubation temperatures and are combined if a protein occured in more than one incubation temperature. CM and CMH are the most common categories. CM and CMH counts are at a similar level despite the difference in detection.

**Fig. S5:**
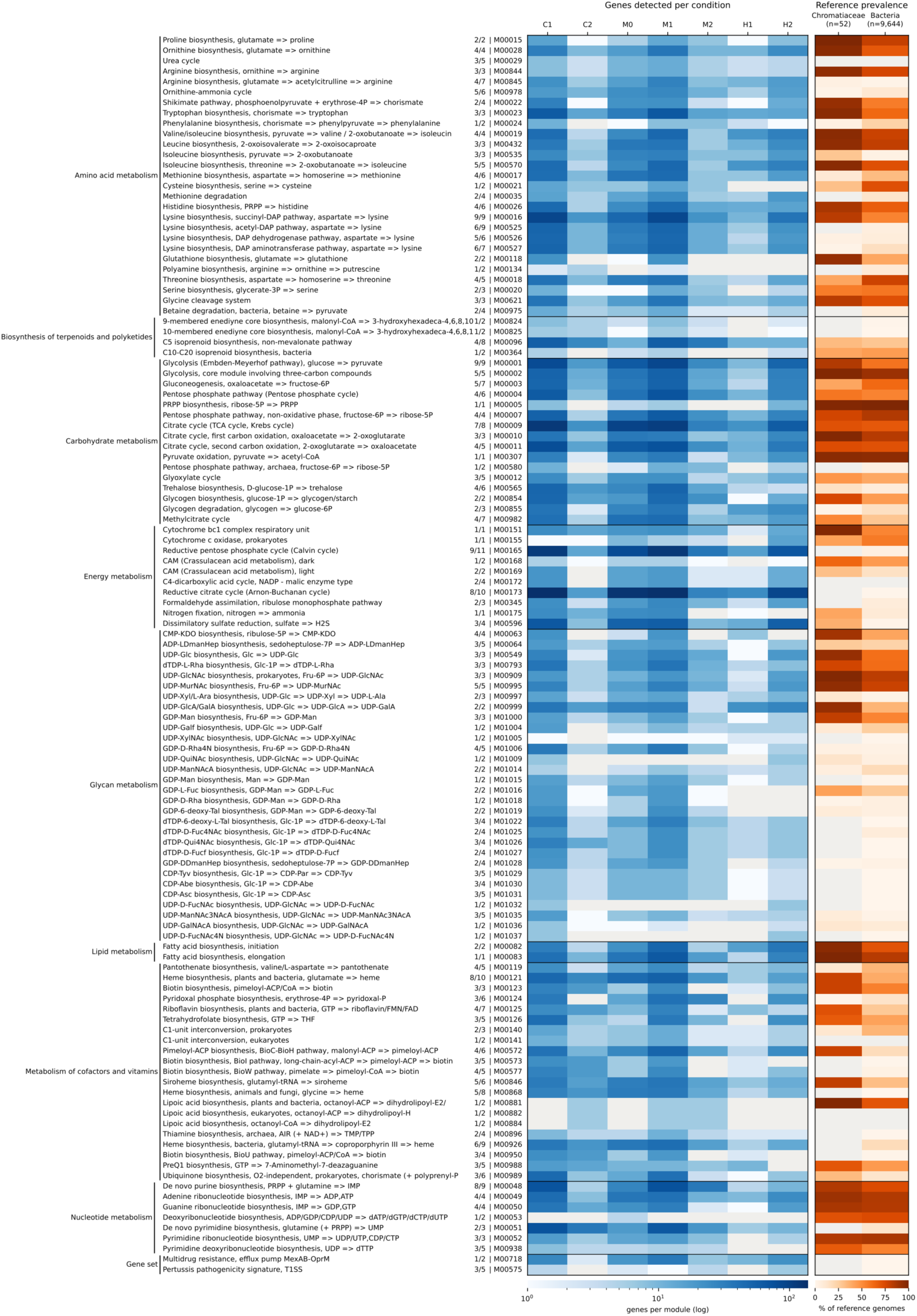
Number of genes identified in proteome per incubation condition that contribute to pathway modules of metabolic relevance. Modules consist of a non-unique set of KIDs, at least 50 % of which are present in the proteome. KIDs were mapped to genes, which therefore are counted multiple times. Occurrence of modules in the reference datasets of all bacteria and chromatiaceae is indicated on the right.

**Fig. S6:**
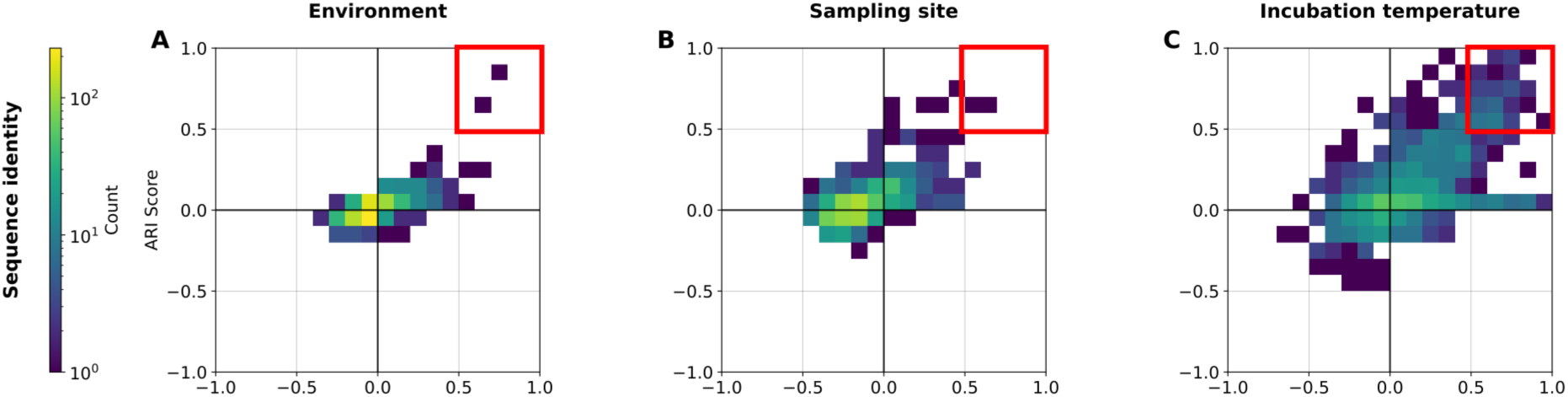
Variant assignment for expressed OGs is independent from environment (A) and shared ancestry (B) with trends preserved from Fig. 1. For incubation temperature (C), the distribution shifted towards positive values, with a 116 OGs surpassing the 0.5 threshold (red box). Clustering based on global alignment distances. Scores for ARI and SIL range between -1 (worse than random), 0 (random) to 1(perfect label matching/geometric separation). Number of OGs per bin is shown in color on a logarithmic scale.

**Tab. S 1:** Genomes used in this study. [genome] identifier used in this study. [Environment] denoting salinity. [Sampling site] denoting type of sampling site and location. Number of contigs in the original assembly [n_contigs], number of nucleotides [n_nuc], as well as stats on maximal and average contig length [max_contig_len, av_contig_len]. Number of contigs after cleaning by CAT [n_contigs_cat]. Number of genes identified by emapper before [n_genes_emapper] and after length-filtering (100-900 residues) [n_genes_len_f]. [Reference] to publication or lab that generated the assembly. Where available, [NCBI] and [IMG ID] identifiers are provided.

| genome | Environ-<br>ment | Sampling site | n_<br>contigs | n_<br>nuc | max_<br>contig_<br>len | av_<br>contig_<br>len | n_<br>contigs_<br>cat | n_<br>genes_<br>emapper | n_<br>genes_<br>len_f | Reference | NCBI | IMG ID |
| --- | --- | --- | --- | --- | --- | --- | --- | --- | --- | --- | --- | --- |
| CAIRSR01_CCP3SC15AN1 | Freshwater | Lake Pavin | 710 | 4,390,319 | 73,281 | 6,183.6 | 577 | 4,166 | 3,181 | <sup>22</sup> | GCA_963575685.1 |  |
| CAIRSR01_CCP3SC1AB1 | Freshwater | Lake Pavin | 261 | 4,287,999 | 101,877 | 16,429.1 | 232 | 3,648 | 2,930 | <sup>22</sup> | GCA_963575655.1 |  |
| CAIRSR01_CCP3SC1AL1 | Freshwater | Lake Pavin | 478 | 3,317,837 | 60,847 | 6,941.1 | 362 | 2,937 | 2,280 | <sup>22</sup> | GCA_963575665.1 |  |
| CAIRSR01_CCP3SC5AM1 | Freshwater | Lake Pavin | 363 | 3,759,029 | 80,373 | 10,355.5 | 295 | 2,832 | 2,364 | <sup>22</sup> | GCA_963575715.1 |  |
| chromatiaceae_CCP4SC76 | Freshwater | Lake Pavin | 825 | 4,082,459 | 57,339 | 4,948.4 | 689 | 3,942 | 2,959 | <sup>22</sup> | GCA_963575725.1 |  |
| bin0 | Freshwater | Lake Stechlin | 2,275 | 7,109,656 | 56,327 | 3,125.1 | 2,058 | 8,598 | 6,093 | <sup>6</sup> |  | 2642422597 |
| bin18 | Freshwater | Lake Stechlin | 8,211 | 9,768,368 | 9,001 | 1,189.7 | 6,676 | 13,211 | 8,307 | <sup>6</sup> |  |  |
| bin2 | Freshwater | Lake Stechlin | 2,604 | 6,095,826 | 37,364 | 2,341.0 | 2,388 | 6,773 | 4,923 | <sup>6</sup> |  |  |
| bin3 | Freshwater | Lake Stechlin | 6,510 | 12,235,064 | 37,126 | 1,879.4 | 5,582 | 15,823 | 10,347 | <sup>6</sup> |  | 2642422598 |
| bin32 | Freshwater | Lake Stechlin | 6,460 | 7,200,153 | 5,007 | 1,114.6 | 5,266 | 11,174 | 6,417 | <sup>6</sup> |  |  |
| bin33 | Freshwater | Lake Stechlin | 7,172 | 6,184,774 | 4,840 | 862.4 | 6,207 | 9,462 | 5,957 | <sup>6</sup> |  |  |
| bin51 | Freshwater | Lake Stechlin | 4,297 | 3,278,107 | 3,657 | 762.9 | 3,810 | 5,884 | 3,615 | <sup>6</sup> |  |  |
| bin6 | Freshwater | Lake Stechlin | 3,933 | 7,745,503 | 18,541 | 1,969.4 | 3,432 | 8,779 | 6,149 | <sup>6</sup> |  |  |
| bin8 | Freshwater | Lake Stechlin | 4,320 | 7,928,622 | 13,013 | 1,835.3 | 3,581 | 8,648 | 5,927 | <sup>6</sup> |  | 2642422596 |
| bin9 | Freshwater | Lake Stechlin | 5,179 | 6,985,974 | 11,863 | 1,348.9 | 4,180 | 9,183 | 5,481 | <sup>6</sup> |  | 2642422595 |
| Cell12 | Freshwater | Lake Stechlin | 816 | 3,637,477 | 71,655 | 4,457.7 | 719 | 3,820 | 2,528 | <sup>6</sup> |  | 2710724217 |
| Cell14 | Freshwater | Lake Stechlin | 10,913 | 10,489,236 | 48,761 | 961.2 | 10,094 | 12,981 | 6,230 | <sup>6</sup> |  |  |
| Cell26 | Freshwater | Lake Stechlin | 2,399 | 3,658,999 | 46,919 | 1,525.2 | 2,144 | 4,904 | 2,780 | <sup>6</sup> |  | 2721755989 |
| Cell3 | Freshwater | Lake Stechlin | 3,478 | 3,893,332 | 21,501 | 1,119.4 | 3,184 | 6,715 | 3,028 | <sup>6</sup> |  | 2711768587 |
| Cell7 | Freshwater | Lake Stechlin | 4,330 | 4,609,114 | 19,095 | 1,064.5 | 3,966 | 7,604 | 3,387 | <sup>6</sup> |  | 2711768588 |
| Cell8 | Freshwater | Lake Stechlin | 1,087 | 4,176,078 | 68,963 | 3,841.8 | 956 | 4,652 | 3,081 | <sup>6</sup> |  | 2710724216 |
| Run1_Cell1 | Freshwater | Lake Stechlin | 1,981 | 1,129,014 | 12,862 | 569.9 | 1,751 | 2,057 | 1,066 | <sup>1</sup> |  |  |
| Run1_Cell2 | Freshwater | Lake Stechlin | 771 | 396,007 | 3,506 | 513.6 | 675 | 834 | 478 | <sup>1</sup> |  |  |
| Run1_Cell3 | Freshwater | Lake Stechlin | 2,050 | 3,882,966 | 97,491 | 1,894.1 | 1,826 | 4,987 | 3,078 | <sup>1</sup> |  |  |
| Run1_Cell4 | Freshwater | Lake Stechlin | 3,064 | 6,781,383 | 103,495 | 2,213.3 | 2,751 | 8,432 | 5,205 | <sup>1</sup> |  |  |
| Run1_Cell5 | Freshwater | Lake Stechlin | 8,417 | 13,223,335 | 51,483 | 1,571.0 | 7,572 | 16,376 | 10,963 | <sup>1</sup> |  |  |
| Run2_Cell1 | Freshwater | Lake Stechlin | 1,339 | 1,358,081 | 17,678 | 1,014.3 | 1,182 | 1,994 | 1,225 | <sup>1</sup> |  |  |
| Run2_Cell12 | Freshwater | Lake Stechlin | 1,714 | 3,191,187 | 45,750 | 1,861.8 | 1,430 | 3,658 | 2,555 | <sup>1</sup> |  |  |
| Run2_Cell13 | Freshwater | Lake Stechlin | 88 | 35,608 | 2,070 | 404.6 | 74 | 62 | 12 | <sup>1</sup> |  |  |
| Run2_Cell15 | Freshwater | Lake Stechlin | 1,363 | 1,078,056 | 14,399 | 790.9 | 1,188 | 1,714 | 927 | <sup>1</sup> |  |  |
| Run2_Cell2 | Freshwater | Lake Stechlin | 2,034 | 3,434,448 | 42,739 | 1,688.5 | 1,682 | 4,057 | 2,739 | <sup>1</sup> |  |  |
| Run2_Cell3 | Freshwater | Lake Stechlin | 244 | 122,194 | 2,805 | 500.8 | 215 | 219 | 91 | <sup>1</sup> |  |  |
| Run2_Cell4 | Freshwater | Lake Stechlin | 1,604 | 2,487,713 | 32,225 | 1,550.9 | 1,394 | 3,123 | 2,078 | <sup>1</sup> |  |  |
| Run2_Cell5 | Freshwater | Lake Stechlin | 4,887 | 3,952,010 | 26,579 | 808.7 | 4,191 | 6,296 | 3,428 | <sup>1</sup> |  |  |
| Run2_Cell7 | Freshwater | Lake Stechlin | 2,105 | 2,393,768 | 19,331 | 1,137.2 | 1,815 | 3,276 | 1,955 | <sup>1</sup> |  |  |
| SC1_sample17_m2A | Freshwater | Lake Stechlin | 1,172 | 343,698 | 2,263 | 293.3 | 1,032 | 1,015 | 196 | <sup>1</sup> |  |  |
| SC1_sample18_m2A | Freshwater | Lake Stechlin | 2,780 | 3,990,981 | 60,797 | 1,435.6 | 2,451 | 4,645 | 2,757 | <sup>1</sup> |  |  |
| SC2_sample19_m2A | Freshwater | Lake Stechlin | 7,470 | 2,693,891 | 8,794 | 360.6 | 6,563 | 6,630 | 1,263 | <sup>1</sup> |  |  |
| SC2_sample20_m2A | Freshwater | Lake Stechlin | 6,281 | 4,853,221 | 58,397 | 772.7 | 5,572 | 7,572 | 3,313 | <sup>1</sup> |  |  |
| SC3_sample21_m2A | Freshwater | Lake Stechlin | 7,215 | 2,598,873 | 10,594 | 360.2 | 6,313 | 6,885 | 1,915 | <sup>1</sup> |  |  |
| SC3_sample22_m2A | Freshwater | Lake Stechlin | 2,529 | 4,096,649 | 136,869 | 1,619.9 | 2,097 | 4,945 | 2,852 | <sup>1</sup> |  |  |
| SC4_sample23_m2A | Freshwater | Lake Stechlin | 2,699 | 4,126,333 | 97,877 | 1,528.8 | 2,439 | 4,860 | 2,813 | <sup>1</sup> |  |  |
| SC4_sample24_m2A | Freshwater | Lake Stechlin | 3,891 | 4,477,605 | 106,888 | 1,150.8 | 3,516 | 5,928 | 3,049 | <sup>1</sup> |  |  |
| SC5_sample25_m2A | Freshwater | Lake Stechlin | 9,237 | 7,250,626 | 68,472 | 785.0 | 8,203 | 11,707 | 5,454 | <sup>1</sup> |  |  |
| SC5_sample26_m2A | Freshwater | Lake Stechlin | 13,727 | 10,896,891 | 52,968 | 793.8 | 12,284 | 16,942 | 8,412 | <sup>1</sup> |  |  |
| A_guadeloupe_bins1 | Saline | Marine | 500 | 3,886,616 | 40,803 | 7,773.2 | 494 | 3,760 | 2,832 | Rémy et al., in prep. |  |  |
| A_guadeloupe_bins3 | Saline | Marine | 383 | 2,764,888 | 41,102 | 7,219.0 | 370 | 2,566 | 2,027 | Rémy et al., in prep. |  |  |
| A_palustre | Saline | Tide pool | 276 | 3,691,389 | 57,832 | 13,374.6 | 276 | 3,397 | 2,652 | <sup>23</sup> | GCA_001044195.1 |  |
| A_sp_WMS1 | Saline | Warm Mineral Springs | 669 | 1,330,408 | 12,602 | 1,988.7 | 632 | 1,660 | 1,189 | <sup>7</sup> | GCA_001276625.1 | 2711768154 |
| A_sp_WMS2 | Saline | Warm Mineral Springs | 777 | 2,849,102 | 47,590 | 3,666.8 | 775 | 3,072 | 2,297 | <sup>7</sup> | GCA_001276605.1 | 8074213743 |
| A_sp_WMS3 | Saline | Warm Mineral Springs | 1,113 | 3,894,228 | 31,412 | 3,498.9 | 1,066 | 4,367 | 2,973 | <sup>7</sup> | GCA_001276615.1 | 2684622566 |

**Tab. S 2:** Functional annotations aggregated per OG. Identifier [og_id] and number of sequences in the OG [n_seqs], most common annotation [consensus_anno] and list of all [annotations] with count.

| og_id | n_seqs | consensus_anno | annotations |
| --- | --- | --- | --- |
| OG0000000 | 5096 | DUF1566 | 'DUF1566', 1181;'Pkinase', 68;'PPC', 13;'Peptidase_S8', 9;'PBP', 6;'Laminin_G_3', 6;'FGE-sulfatase', 4;'Beta_helix', 4;'ThiS', 2;'He_PIG', 2;'Cellulase', 1;'RHS_repeat', 1;'fn3', 1;'DUF3631', 1;'Fibrinogen_C', 1;'PhoPQ_related', 1 |
| OG0000001 | 2452 | FGE-sulfatase | 'FGE-sulfatase', 448;'Peptidase_C14', 20;'DUF1566', 8;'Pkinase', 7;'PEGA', 5;'DUF4198', 2;'DinB_2', 1;'RVT_1', 1 |
| OG0000002 | 1629 | N6_Mtase | 'N6_Mtase', 102 |
| ... | ... | ... | ... |

**Tab. S 3:**
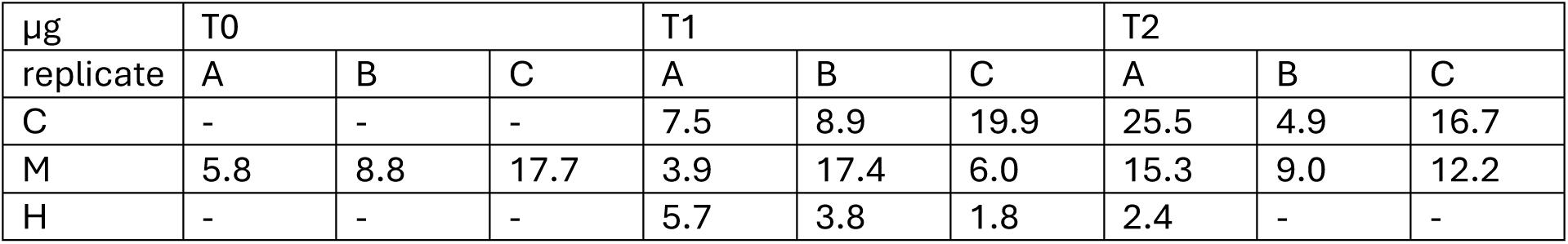
Protein amounts per sample after stacking gel in µg. Timepoints 1 day [T0], 2 weeks [T1] and 4 weeks [T2] after sampling. Incubation temperatures 7 °C [C], room temperature [M] and 30 °C [H]. Replicates denoted A-C. No sample provided is indicated with ‘–‘.

**Tab. S 4:**
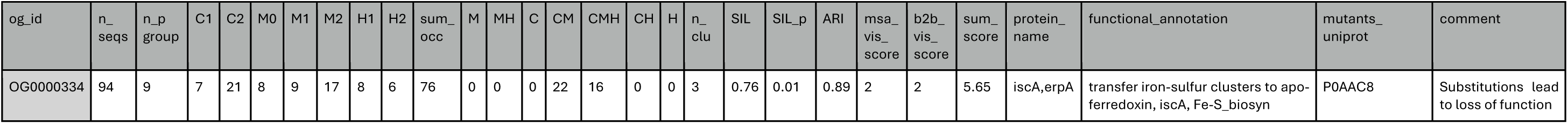
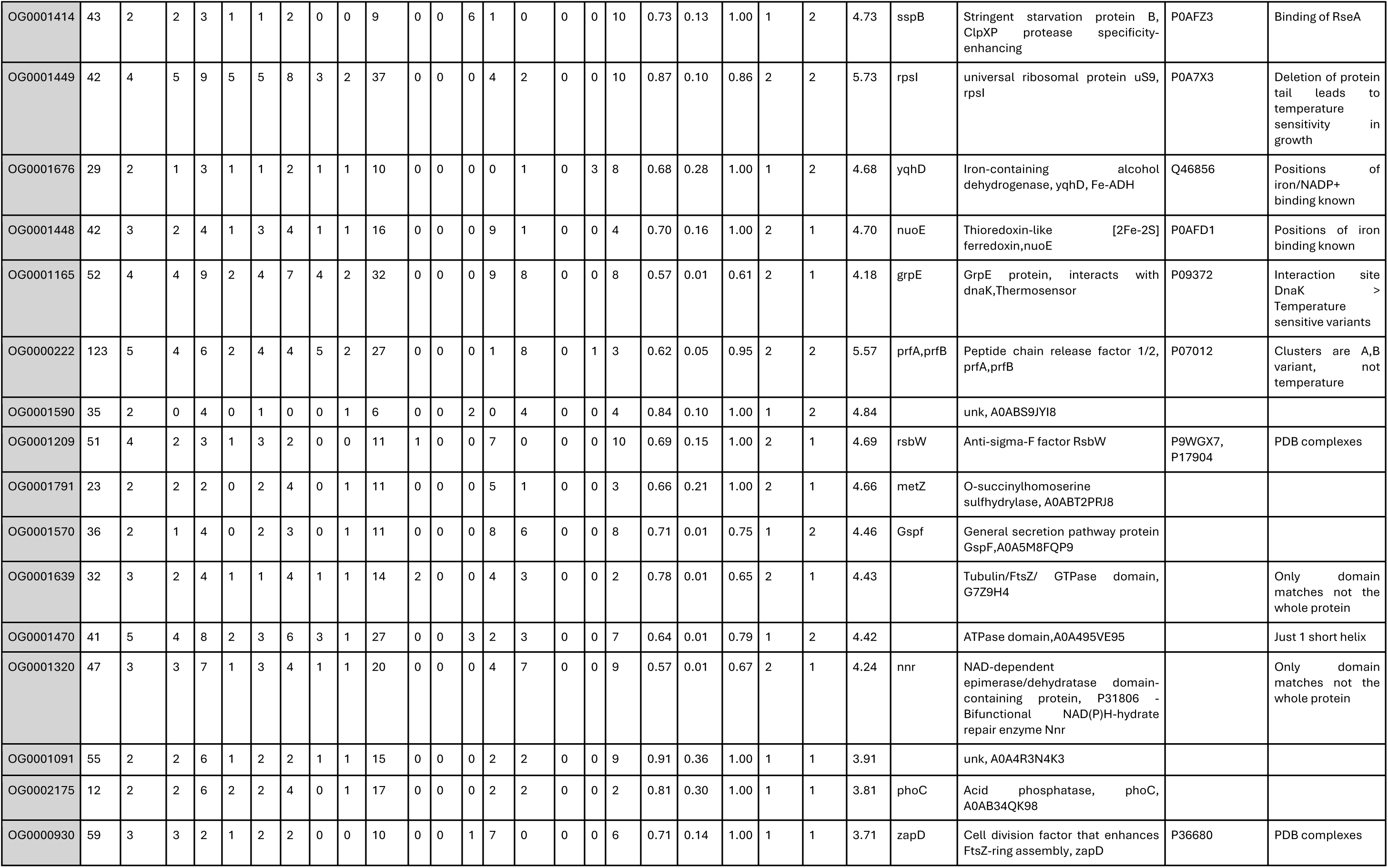
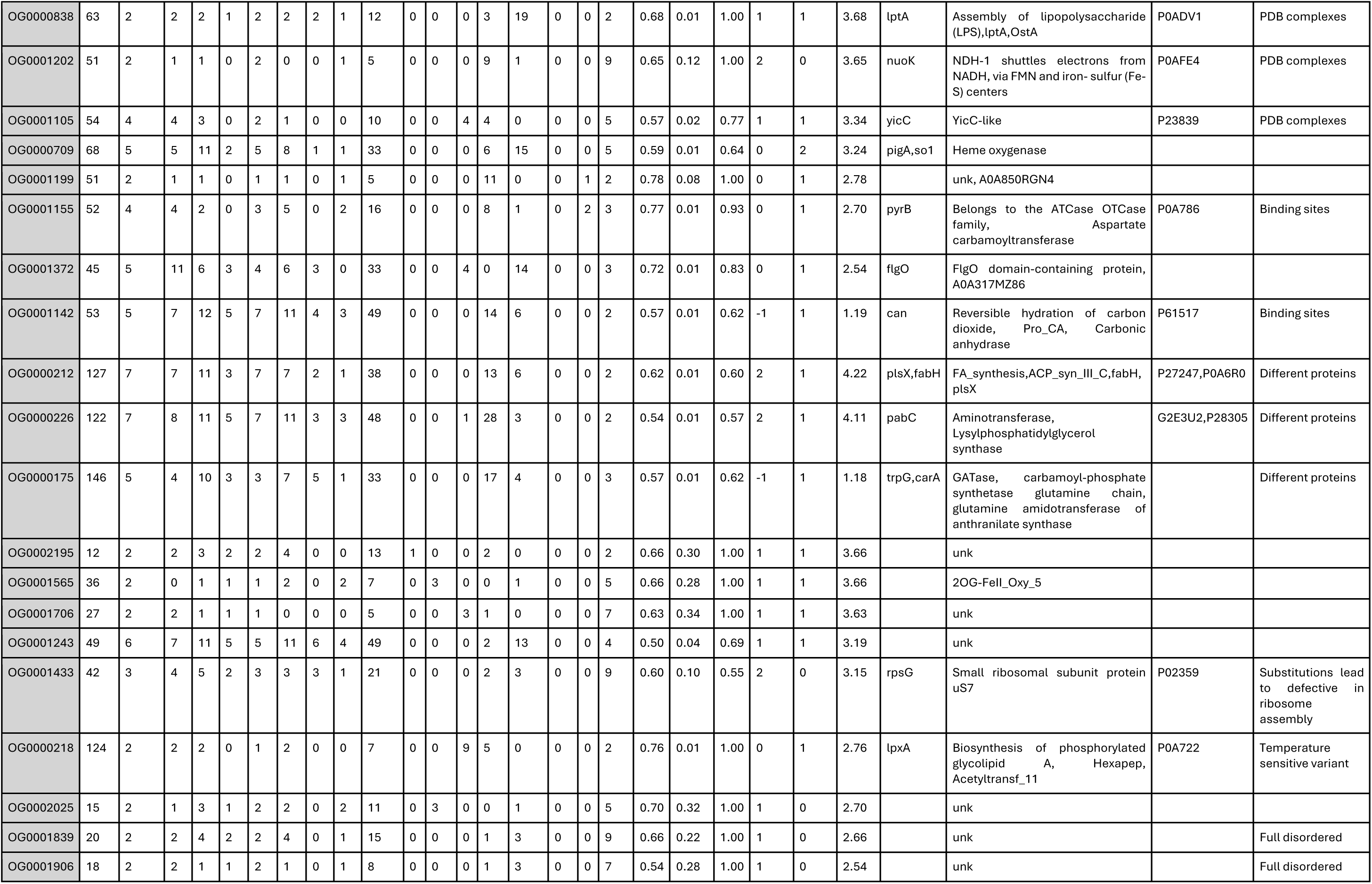
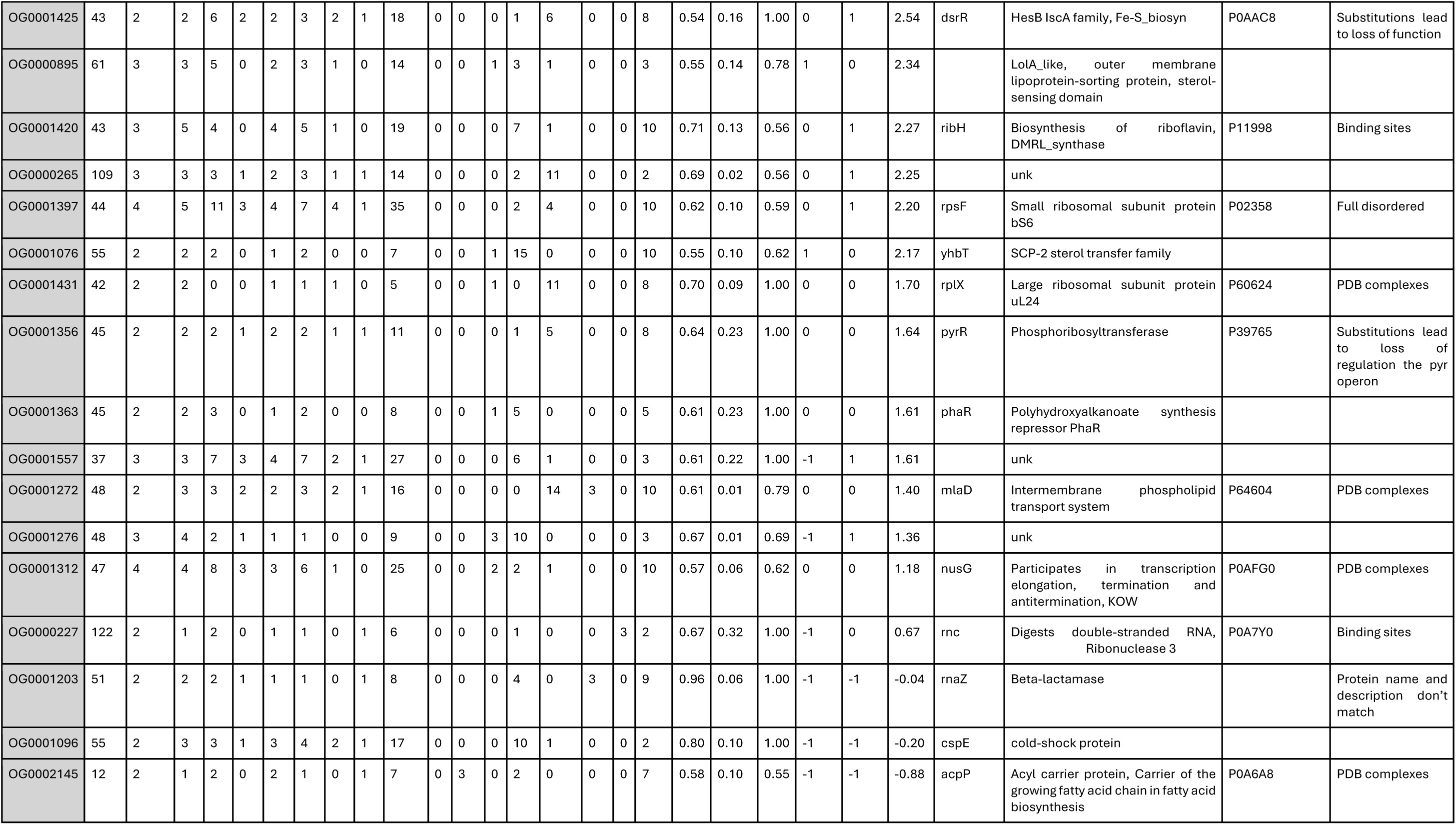
Selected OGs scoring sheet. Identifier [og_id] and number of sequences (seq) in the theoretical [n_seqs] versus experimental [n_pgroup] proteome (one pgroup can map to multiple seqs). Occurrence per temperature/timepoint for all replicates [C,M,H,0,1,2] and total [sum_occ] in MS-proteomics (each seq can appear multiple times). Counts for incubation temperature categories [C,CM,CMH,CH,H] in MS-proteomics (each seq can appear once). Number of clusters determined by k-Means [n_clu]. SIL score and its p-value for to assess temperature category clustering [SIL,SIL_p]. [ARI] score assessing agreement between temperature category and k-Means clustering. [msa_vis_score] visual score of MSA quality and variation. [b2b_vis_score] visual score of difference between backbone dynamics per k-Means cluster. [sum_score] ranking OG potential by adding SIL,ARI and the vis_scores. Consensus functional annotation from emapper and BLAST [protein_name, functional_annotation]. UniProt ID of well annotated bacterial ortholog [mutants_uniprot] identified by protein name and verified by predicted 3D structure. [Comment] contains interaction/mutation that may be interpreted via biophysical propensities.

